# Loss of NKX2-1 predisposes thyroid to neoplasm development through regulation of oxidative stress

**DOI:** 10.64898/2026.08.26.746581

**Authors:** Yo-Taro Shirai, Jerrold M. Ward, Yoshinori Takizawa, Huaitian Liu, Masaaki Miyakoshi, Manabu Iwadate, Tsubasa Murata, Suguru Hayase, Shigetoshi Yokoyama, Shogo Ehata, Shioko Kimura

## Abstract

Many factors including ionizing radiation and iodine deficiency are known to increase thyroid carcinogenesis risk. Our dataset analysis of The Cancer Genome Atlas (TCGA) showed that lower mRNA expression of *NK2 homeobox 1* (*NKX2-1*) transcription factor, a master regulator of genesis, homeostasis, and function of thyroid, is linked to poor prognosis of papillary thyroid cancer patients. Here we provide the findings that thyroid-specific *Nkx2-1* conditional knockout (*Nkx2-1*^ΔT^) mice develop thyroid adenoma and carcinoma in higher frequency with combined exposure to radiation and iodine deficiency than control *Nkx2-1*^fl/fl^ mice. Iodine deficiency caused oxidative stress, which subsequently resulted in DNA damage, leading to transformation of thyroid follicular cells. RNA-seq gene set enrichment analysis indicated higher production of reactive oxygen species (ROS) in the thyroids of *Nkx2-1*^ΔT^ as compared to *Nkx2-1*^fl/fl^ mice with combined exposure to radiation and iodine deficiency. This was accompanied by a feedback induction of SOD3 (superoxide dismutase 3) and GPX2 (glutathione peroxidase 2). These antioxidants were naturally expressed at higher levels in the thyroids of *Nkx2-1*^ΔT^ than *Nkx2-1*^fl/fl^ mice without iodine deficiency or radiation. *Nkx2-1*^ΔT^ thyroids exhibited abnormal follicle architecture and up-regulation of *Acox2* (encoding acyl-CoA oxidase 2), which produces hydrogen peroxide. These results suggest that loss of NKX2-1 may contribute to excess ROS production, which elevates basal oxidative stress resulting in the promotion of ROS-induced carcinogenesis. We propose a role for NKX2-1 as a regulator of ROS production homeostasis in the thyroid. Its disturbance would dispose thyroid follicular cells more vulnerable to the ROS-producing carcinogens.

## Introduction

Thyroid is an endocrine organ which possesses a unique metabolism system utilizing iodide and hydrogen peroxide (H_2_O_2_) generated by dual oxidase (DUOX) to produce thyroid hormone [1]. Ionizing radiation emitted from radioiodine is a well-known risk factor for thyroid carcinogenesis through DNA damage and genomic instability as seen in the relationship between the disastrous nuclear power plant accident in Chernobyl and thyroid cancer patients at young age [2, 3]. In addition, iodine deficiency is responsible for high incidence of thyroid disorders in Europe [4] and was reported to induce tumor in rat thyroids [5]. These factors are thought to play important roles in thyroid carcinogenesis most likely related to the thyroid-specific metabolism system during thyroid hormone synthesis.

Thyroid cancer originates from thyroid follicular cells except medullary carcinoma and morphologic types include papillary thyroid carcinoma (PTC), follicular thyroid carcinoma (FTC), oncocytic thyroid carcinoma (OC, also known as Hürthle cell carcinoma), poorly differentiated thyroid carcinoma (PDTC), and anaplastic thyroid carcinoma (ATC) [6]. PTC, the most common histopathological type of thyroid cancer (up to ∼90%), FTC with the second highest prevalence (up to ∼10%) and OC are differentiated thyroid cancers with good prognosis [6, 7]. The fact encompassing clinical significance is that PDTC and ATC with more aggressive invasion and poor prognosis arise from PTC and FTC through de-differentiation. It should be noted that FTC is thought to develop from thyroid adenoma as proposed in a multi-step thyroid carcinogenesis model [7], thus, it is essential to fully clarify the thyroid carcinogenesis mechanisms including thyroid adenoma.

NK2 homeobox 1 (NKX2-1, also known as thyroid transcription factor-1; TTF-1), is a homeodomain transcription factor known as a master regulator of thyroid development and differentiation as well as lung and some parts of brain [8–11]. NKX2-1 is also crucial for the maintenance of organized thyroid architecture, thyroid function and expression of thyroid-specific genes involved in hormone synthesis [10, 12]. Relationship between NKX2-1 and lung cancer has been intensely investigated, and dual, oncogenic and tumor suppressive roles of NKX2-1 were reported [13]. On the other hand, the role of NKX2-1 in thyroid cancer is largely unknown. Our previous study demonstrated the increased incidence of thyroid adenoma development induced by a combination of genotoxic chemical mutagen N-bis(2-hydroxypropyl)-nitrosamine (DHPN) and goitrogen sulfadimethoxine (SDM) in thyroid-specific *Nkx2-1* conditional knockout (*Nkx2-1*^ΔT^) as compared with control *Nkx2-1*^fl/fl^ mice [14]. However, chemical-induced thyroid cancer is very limited and may have specific mechanisms, thus the effects of loss of NKX2-1 on other common thyroid-specific cancer risk factors need to be clarified.

In this study, a combination of radiation and iodine deficiency together with the loss of *Nkx2-1* which presented poor prognosis of PTC patients on the dataset of The Cancer Genome Atlas (TCGA) was chosen to study whether and/or how they affect thyroid carcinogenesis in mice. Intrinsically increased reactive oxygen species (ROS) production demonstrated with enhanced expression of antioxidants was found in *Nkx2-1*^ΔT^ mouse thyroids as compared with normal *Nkx2-1*^fl/fl^ thyroids. This was further enhanced by iodine deficiency and radiation, which led to enhanced DNA damage and increased incidence of carcinogenesis. In this study, we offer an insight into the relationship between NKX2-1 and thyroid carcinogenesis; loss of NKX2-1 is an important risk factor for radiation and iodine deficiency-induced thyroid carcinogenesis through regulation of ROS.

## Results

### Poor prognosis of PTC patients with low expression of *NKX2-1*

The clinical outcome of *NKX2-1* expression in thyroid cancer patients was first examined using TCGA database, focusing on the most common type of thyroid cancer, PTC, including follicular patterned and tall cell patterned (Figures 1A and S1). The patients with low *NKX2-1* expression had significantly poorer disease-specific prognosis than the patients with high and intermediate *NKX2-1* expression (Figure 1A). This finding suggests that NKX2-1 may work as a tumor suppressor in thyroid cancer.

**Figure 1.**
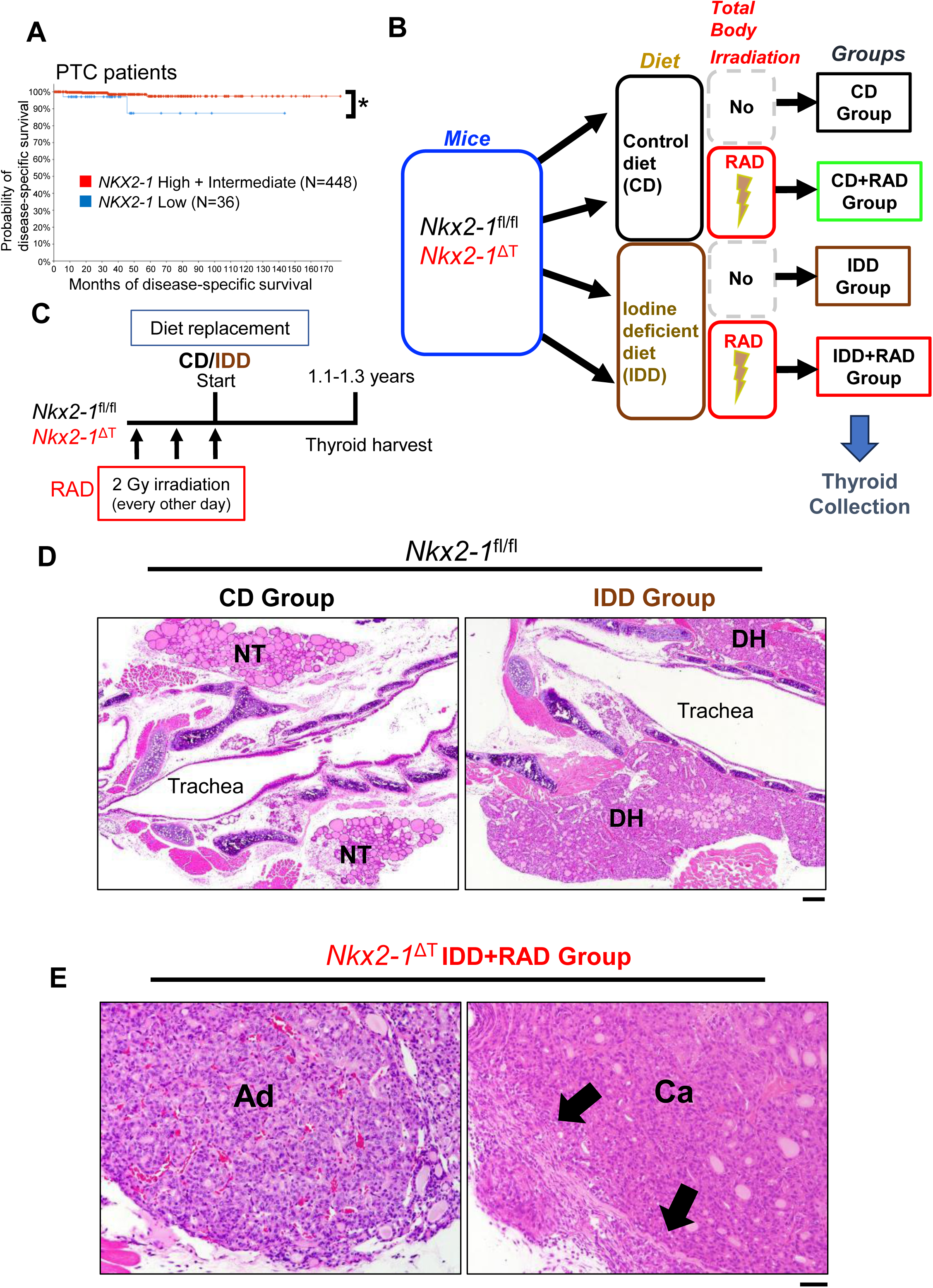
Thyroid carcinogenesis induced by radiation and iodine deficiency in *Nkx2-1* thyroid-specific conditional knockout mice. (A) Disease-specific survival of papillary thyroid carcinoma (PTC) patients based on *NKX2-1* expression obtained from Thyroid Carcinoma TCGA PanCancer Atlas dataset. *p<0.05 by Logrank test. (B) Schematic representation of grouping for mouse thyroid carcinogenesis study of iodine deficiency and radiation using *Nkx2-1*^fl/fl^ and *Nkx2-1*^ΔT^ mice (CD, CD+RAD, IDD, IDD+RAD). (C) Schematic representation of time course of the study described in (B). (D and E) Representative H&E (Hematoxylin and Eosin)-stained mouse thyroid histology pictures observed in the study. Normal thyroid gland under CD and thyroid diffuse hyperplasia (goiter) induced by iodine deficiency in *Nkx2-1*^fl/fl^ mice (D). Follicular adenoma and carcinoma in IDD+RAD-treated *Nkx2-1*^ΔT^ mouse thyroids (E). Capsule invasion indicated with arrows. Magnification: X4 objective (D), X20 objective (E). NT, normal thyroid; DH, diffuse hyperplasia; Ad, adenoma; Ca, carcinoma. Bar, 200 μm.(D), 50 μm. (E).

### Thyroid-specific *Nkx2-1* conditional knockout mice develop higher number of thyroid adenomas with combined exposure to radiation and iodine deficiency

In order to examine the effects of gamma-ray irradiation and iodine deficiency in combination with the loss of *Nkx2-1* on thyroid carcinogenesis, male *Nkx2-1*^fl/fl^ control mice and thyroid-specific *Nkx2-1* conditional knockout mice *Nkx2-1*^fl/fl^;*TP*O-Cre (*Nkx2-1*^ΔT^) [12, 15] were randomly allotted to four groups described as follows: CD group (Control diet, no Radiation); CD+RAD group (Control diet+Radiation; total body gamma-ray irradiation (2 Gy x 3)); IDD group (Iodine deficient diet, no Radiation); IDD+RAD group (Iodine deficient diet+Radiation; Figure. 1B, see details in Materials Methods). Over one year after the diet replacement at the age of 5.0 to 8.0 weeks old, thyroids were harvested for histological examination (Figure 1C). First, the effect of iodine deficiency was confirmed with *Nkx2-1*^fl/fl^ mice, as all the thyroids in CD group showed normal follicles without any abnormal lesions as expected, and most IDD-treated thyroids displayed diffuse hyperplasia (goiter) as reported in mice [16] and rats [17] (Figure 1D; Table S1). While radiation induced focal hyperplasia in some *Nkx2-1*^fl/fl^ mice, it did not induce adenoma or carcinoma (Tables 1 and S1). Adenoma formation was observed in IDD and IDD+RAD groups, but not CD+RAD group (Table 1), suggesting that iodine deficiency may work as an inducer of adenoma formation, but not radiation in this model. Carcinoma developed only in IDD+RAD group, suggesting a possibility that radiation further promotes IDD-induced genomic instability, leading to carcinoma development as proposed in a multi-step carcinogenesis model [7] (Table 1).

**Table 1.** Summary of adenoma and carcinoma incidence in *Nkx2-1*^fl/fl^ and *Nkx2-1*^ΔT^ mouse thyroids with combined exposure to radiation and iodine deficiency over one year since diet replacement.

| Genotype | Diet/<br>Radiation | Adenoma (%)<br>(mouse number) | Carcinoma (%)<br>(mouse number) | Total mouse<br>number (N) |
| --- | --- | --- | --- | --- |
| *Nkx2-1<sup>fl/fl</sup>* | CD | 0<br>(0) | 0<br>(0) | 21 |
| *Nkx2-1<sup>ΔT</sup>* |  | 0<br>(0) | 0<br>(0) | 23 |
| *Nkx2-1<sup>fl/fl</sup>* | CD+RAD | 0<br>(0) | 0<br>(0) | 22 |
| *Nkx2-1<sup>ΔT</sup>* |  | 0<br>(0) | 0<br>(0) | 28 |
| *Nkx2-1<sup>fl/fl</sup>* | IDD | 6<br>(1) | 0<br>(0) | 18 |
| *Nkx2-1<sup>ΔT</sup>* |  | 23<br>(7) | 0<br>(0) | 30 |
| *Nkx2-1<sup>fl/fl</sup>* | IDD+RAD | 15<br>(4) | 4<br>(1) | 27 |
| *Nkx2-1<sup>ΔT</sup>* |  | 50**<br>#(17) | 9<br>(3) | 35<br>(34 for adenoma) |

Next, frequency of adenoma and carcinoma formation between *Nkx2-1*^fl/fl^ and *Nkx2-1*^ΔT^ mice was compared. Significantly higher incidence of adenoma formation was obtained in *Nkx2-1*^ΔT^ mice in IDD+RAD group in addition to higher carcinoma development incidence although without statistical significance (Figure 1E; Table 1). The results were reproduced using another set of mice (Table S2), where carcinoma incidence was very low or none without homologous deletion of *Nkx2-1*. These results suggest that the loss of *Nkx2-1* may promote IDD+RAD-induced thyroid carcinogenesis.

### Iodine deficiency generates ROS and DNA damage in thyroids

Ionizing radiation is known to directly damage DNA and also generate free radicals including ROS which eventually cause DNA damages as indirect toxic effects on organs [18, 19]. In addition, excess H_2_O_2_, another ROS, is thought to be produced by iodine deficiency due to imbalance of iodide oxidization for coupling with thyroglobulin [20]. Previous study showed that iodine deficiency induces DNA damage with enhanced activity of superoxide dismutase (SOD) [21] in rat thyroids and increases expression of *Sod3* in mouse thyroids [22]. Based on these, we hypothesized that iodine deficiency and radiation independently generate ROS and have additive effects on thyroid carcinogenesis (Figure 2A). Since radiation alone did not induce adenoma or carcinoma development in the mouse thyroids (Table 1), DNA damage by iodine deficiency with/without radiation for 13 weeks was next assessed in *Nkx2-1*^fl/fl^ mouse thyroids (Figure 2B) by examination of phosphorylated form of Histone 2A variant X (γH2AX), a sensitive marker for DNA double-strand break damage and repair [19, 23]. γH2AX was clearly induced in both IDD and IDD+RAD groups although augmentation by radiation was not detected (Figure 2C), suggesting that iodine deficiency may potently induce strong DNA damage in mouse thyroids as reported [22].

**Figure 2.**
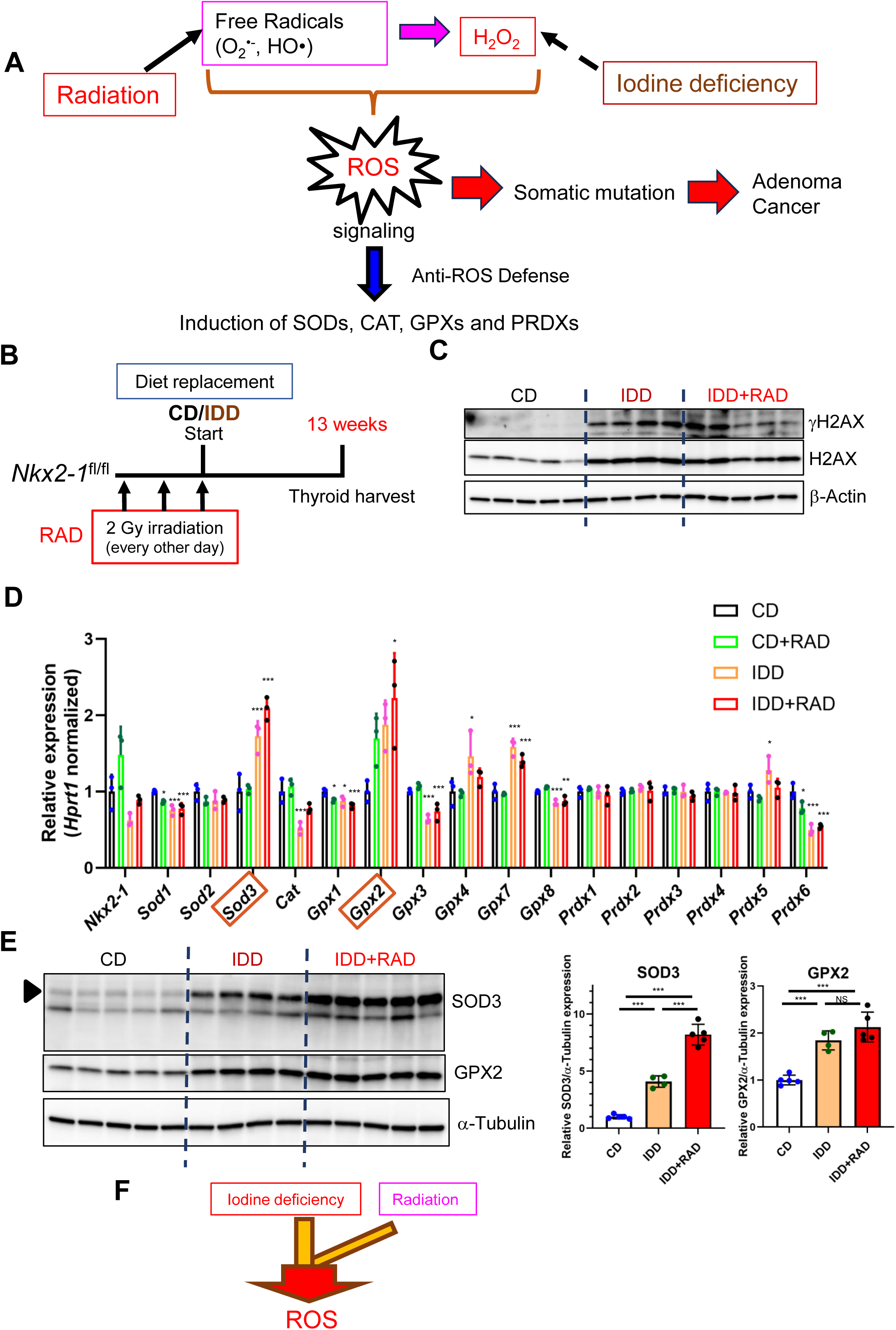
Iodine deficiency causes DNA damage and induces expression of SOD3 and GPX2 antioxidants in mouse thyroids. (A) Schematic representation of hypothesis that radiation and iodine deficiency independently produce ROS. Produced ROS functions as a signaling molecule and results in induction of antioxidants as a defense system against it. Overproduced ROS causes DNA damage and subsequent DNA mutations, which eventually lead to neoplasm development including adenoma and cancer. (B) Schematic representation of time course of molecular profile analysis using *Nkx2-1*^fl/fl^ mouse thyroids with or without IDD/RAD. Thyroids were harvested at 13 weeks after diet replacement (CD, CD+RAD, IDD, IDD+RAD groups). (C and E) Immunoblotting of mouse thyroid lysates in CD, IDD, and IDD+RAD groups with indicated antibodies (γH2AX, #9718). The same lysates were used for (C) and (E). β-Actin (C) and α-Tubulin (E) were used as loading controls, respectively. Quantification of SOD3 or GPX2/α-Tubulin (Right, E). ***p<0.005 by Tukey’s multiple comparison tests. NS, not significant. (D) qRT-PCR for *Nkx2-1* and indicated representative antioxidant gene mRNAs using cDNAs obtained from *Nkx2-1*^fl/fl^ mouse thyroids in CD, CD+RAD, IDD, and IDD+RAD groups. Data are presented as mean ± SD from individual mouse thyroid samples (N=3 for each group). For each gene, the mean value of CD group samples was set as 1. *p<0.05, **p<0.01, ***p<0.005 for comparison against CD group samples by Dunnett’s multiple comparison tests. Significantly up-regulated genes with >2 fold increase in IDD+RAD group are marked with red rectangles. (F) Schematic model of ROS production by iodine deficiency and radiation. Iodine deficiency dominantly produces ROS and radiation further enhances ROS production in our study.

ROS functions as a signaling molecule and is known to induce expression of antioxidants including SODs, catalase (CAT), glutathione peroxidases (GPXs), peroxiredoxins (PRDXs) as a feedback defense [24–27] (Figure 2A). The highest mRNA expression of *Sod3* and *Gpx2* with >2 fold induction was found among the direct-acting antioxidants examined in IDD+RAD, followed by IDD as compared with CD group (Figure 2D). The highest increase in SOD3 and GPX2 expression by IDD+RAD was also confirmed at protein level, with lower induction level by IDD alone at least regarding SOD3 (Figure 2E). These results suggest that iodine deficiency may generate ROS and DNA damage with further enhancement by ionizing radiation in the thyroid (Figure 2F).

### Enrichment of oxidative stress pathways and antioxidant molecules in *Nkx2-1*^ΔT^ thyroids

RNA-seq was next performed using mouse thyroids collected after 4 weeks of IDD+RAD treatment to reveal the molecular profile differences between *Nkx2-1*^fl/fl^ and *Nkx2-1*^ΔT^ thyroids (Figure 3A). Gene set enrichment analysis (GSEA) [28] revealed enrichment of oxidative stress pathways (Figure 3B; Tables S3 and S4), suggesting a possibility that ROS production by IDD+RAD was more severe in *Nkx2-1*^ΔT^ thyroids. In accordance with this, increased expression of glutathione S-transferase (GST) [29], the enzyme required for transfer of glutathione (GSH), was seen in *Nkx2-1*^ΔT^ thyroids, and the increase was confirmed by qRT-PCR (Figure S2). Among the direct-acting antioxidants, *Sod3* and *Gpx2* were clearly up-regulated (Figure 3C), as confirmed at mRNA and protein levels (Figures 3D and 3E). Taken together with the observation of the highest induction of SOD3 and GPX2 by IDD+RAD (Figures 2D and 2E), the finding here also supports higher ROS production in *Nkx2-1*^ΔT^ mouse thyroids.

**Figure 3.**
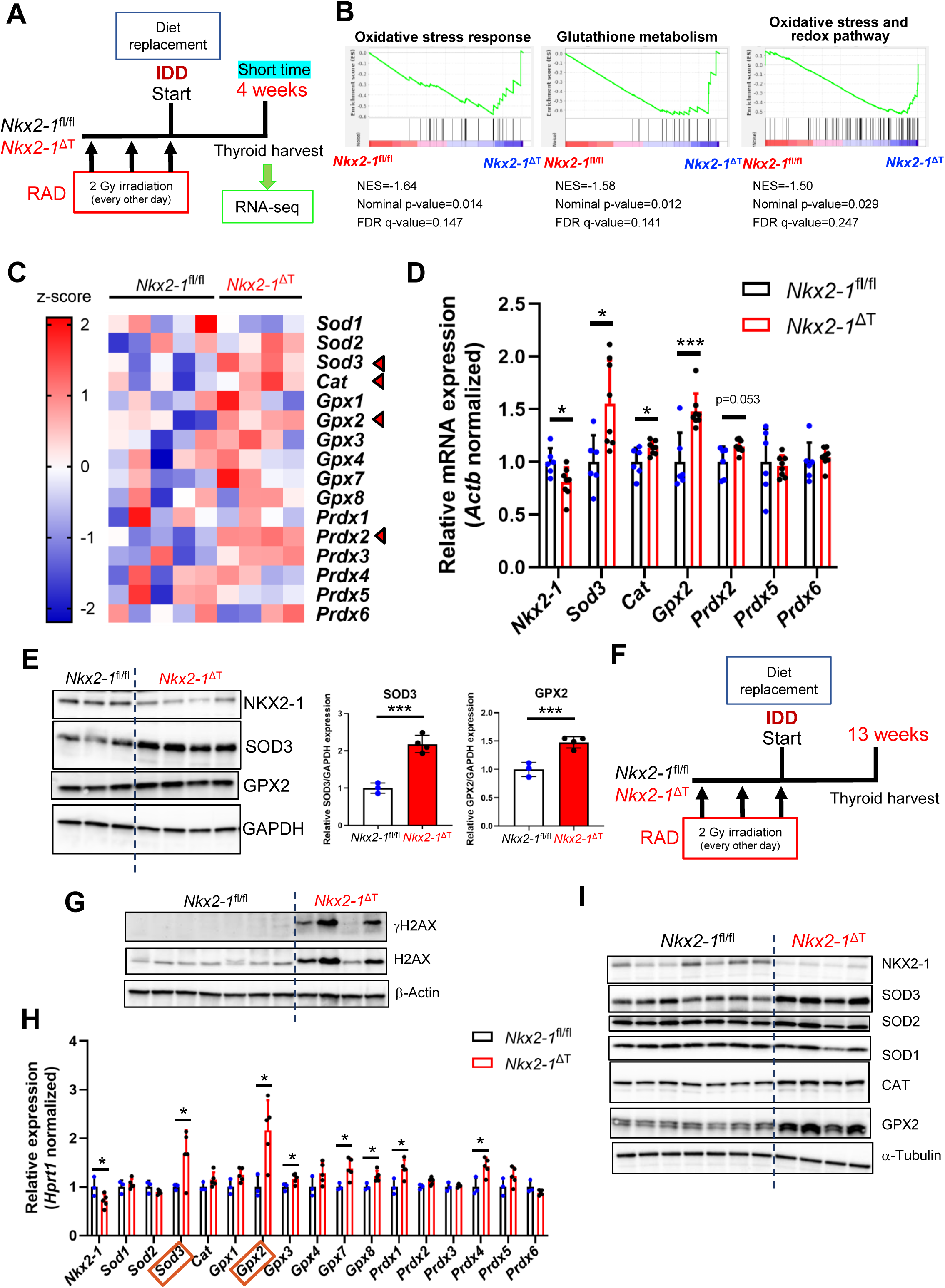
ROS production is enhanced in *Nkx2-1*^ΔT^ mouse thyroids under a combination of iodine deficiency and radiation. (A) Schematic representation of short-time course experiments for RNA-seq of IDD+RAD-treated *Nkx2-1*^fl/fl^ and *Nkx2-1*^ΔT^ mouse thyroids. (B and C) RNA-seq using RNAs obtained from thyroids of *Nkx2-1*^fl/fl^ (N=5) and *Nkx2-1*^ΔT^ (N=4). Gene set enrichment analysis (GSEA) was performed and representative enriched datasets regarding oxidative stress in *Nkx2-1*^ΔT^ (WikiPathway subset) are shown in (B). Nominal p-values and FDR q-values as indicated. NES, normalized enrichment score. See details in Tables S3 and S4. Heatmap of representative antioxidant genes is shown in (C). Significantly up-regulated genes in *Nkx2-1*^ΔT^ are indicated with red arrows. (D) qRT-PCR for *Nkx2-1* and indicated representative antioxidant gene mRNAs using cDNAs from *Nkx2-1*^fl/fl^ (N=6) and *Nkx2-1*^ΔT^ (N=8) mouse thyroids collected at 4 weeks after diet replacement as described in (A). RNA samples used for this analysis include those used for RNA-seq. Data are presented as mean ± SD from individual mouse thyroid samples. For each gene, the mean value of *Nkx2-1*^fl/fl^ mouse samples was set as 1. *p<0.05, ***p<0.005 by Student’s *t*-tests. See Materials & Methods for *Nkx2-1* deletion in *Nkx2-1*^ΔT^ thyroid follicular cells. (E) Immunoblotting of *Nkx2-1*^fl/fl^ (N=3) and *Nkx2-1*^ΔT^ (N=4) mouse thyroid lysates obtained in the same time course as described in (A) with indicated antibodies (Left). Quantification of SOD3 or GPX2/GAPDH (Right). Mean value of *Nkx2-1*^fl/fl^ mouse samples was set as 1. ***p<0.005 by Student’s *t*-tests. (F) Schematic representation of 13 week-time course experiments for *Nkx2-1*^fl/fl^ and *Nkx2-1*^ΔT^ mouse thyroids after diet replacement in IDD+RAD group. (G and I) Immunoblotting of *Nkx2-1*^fl/fl^ (N=7) and *Nkx2-1*^ΔT^ (N=4) mouse thyroid lysates as described in (F) with indicated antibodies (γH2AX, #9718; CAT, #14097). (H) qRT-PCR for *Nkx2-1* and indicated representative antioxidant gene mRNAs using cDNAs from *Nkx2-1*^fl/fl^ (N=3) and *Nkx2-1*^ΔT^ (N=5) mouse thyroids as described in (F). Data are presented as mean ± SD from individual mouse thyroid samples. For each gene, the mean value of *Nkx2-1*^fl/fl^ mouse samples was set as 1. *p<0.05 by Student’s *t*-tests or Welch’s *t*-test. Significantly up-regulated genes with >1.5 fold increase in *Nkx2-1*^ΔT^ mouse thyroids are marked with red rectangles.

### Higher extent of DNA damage by a combination of iodine deficiency and radiation in *Nkx2-1*^ΔT^ mouse thyroids

In order to confirm the finding described above, γH2AX was also examined in *Nkx2-1*^fl/fl^ and *Nkx2-1*^ΔT^ thyroids with exposure to radiation and iodine deficiency for 13 weeks as shown in the scheme (Figure 3F). Clearly higher γH2AX expression was detected in most *Nkx2-1*^ΔT^ thyroids (Figure 3G), suggesting that DNA damage caused by IDD+RAD is stronger in *Nkx2-1*^ΔT^ thyroids. Expression of many direct-acting antioxidant gene mRNAs was up-regulated in *Nkx2-1*^ΔT^ thyroids (Figure 3H). It should be noted that only *Sod3* and *Gpx2* showed significant increase with >1.5 fold at mRNA level among the direct-acting antioxidant genes examined (Figure 3H). Expression levels of these proteins were also clearly increased in *Nkx2-1*^ΔT^ thyroids (Figure 3I). Further, increased expression of γH2AX, SOD3 and GPX2 in *Nkx2-1*^ΔT^ thyroids was also confirmed at 21 weeks after diet replacement (Figure S3). These results suggest that higher ROS production causing higher degree of DNA damage took place in the thyroids of *Nkx2-1*^ΔT^ than *Nkx2-1*^fl/fl^ mice in IDD+RAD group.

### Higher oxidative stress with up-regulation of thyroid hormone synthesis-related gene expression in *Nkx2-1*^ΔT^ thyroids

Next, experiments were carried out to determine whether the effect of loss of NKX2-1 on oxidative stress is specifically observed only in the presence of ROS-producing exogenous stimuli. Expression levels of antioxidants between *Nkx2-1*^fl/fl^ and *Nkx2-1*^ΔT^ thyroids were compared 13 weeks after the start of CD without any additional ROS-producing factors (Figure 4A). Interestingly, clearly higher expression of *Sod3* and *Gpx2* (>2 fold) was again observed in CD group of *Nkx2-1*^ΔT^ thyroids (Figure 4B). These results suggest a possibility that *Nkx2-1*^ΔT^ thyroids possess elevated basal oxidative stress which makes thyroid follicular cells more vulnerable to ROS-induced genomic instability, and as a result develop adenoma and carcinoma in higher frequency under IDD+RAD.

**Figure 4.**
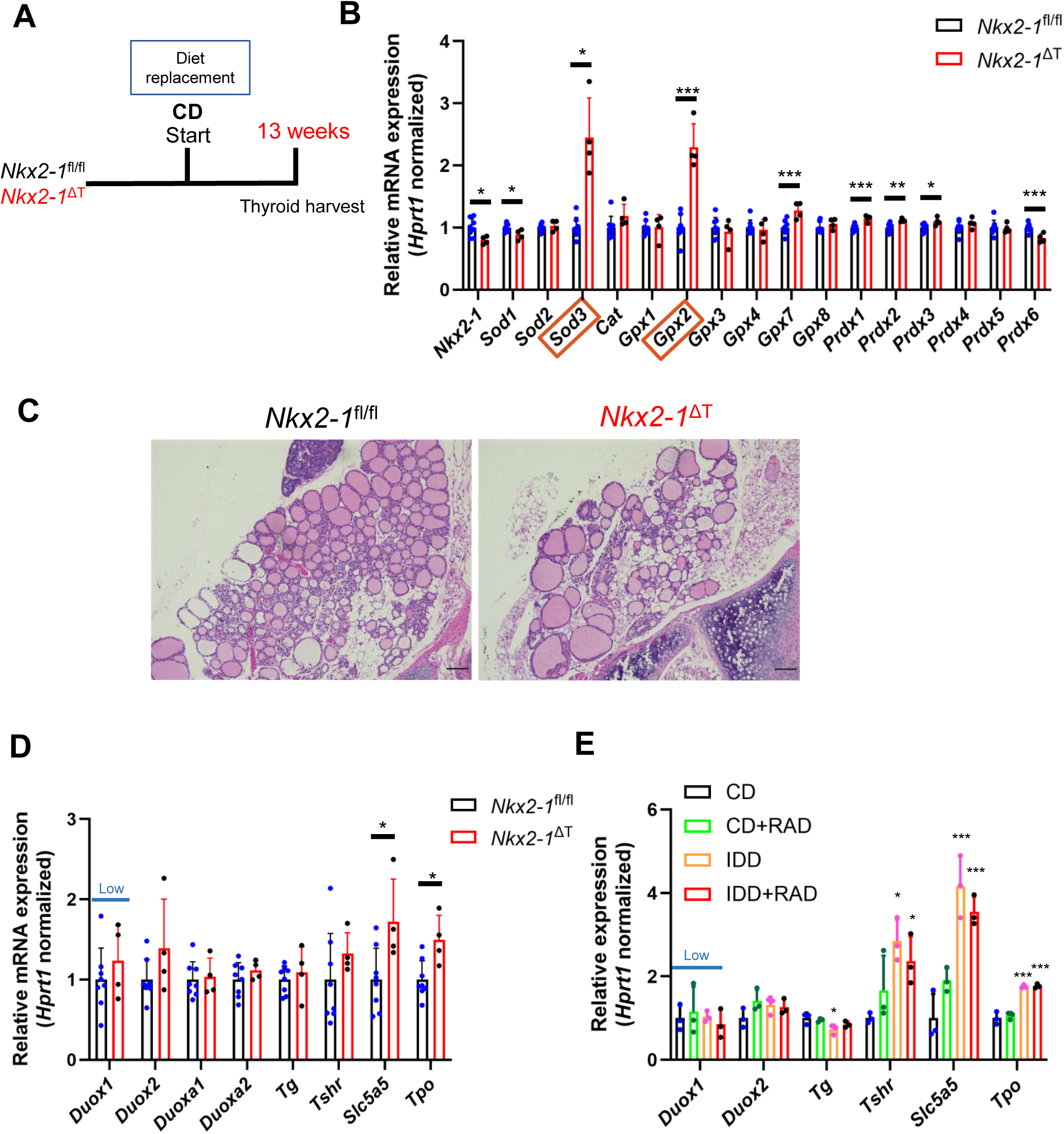
Up-regulation of *Slc5a5* and *Tpo* expression in *Nkx2-1*^ΔT^ mouse thyroids. (A) Schematic representation of 13 week-time course experiments for *Nkx2-1*^fl/fl^ and *Nkx2-1*^ΔT^ mouse thyroids after diet replacement with CD. (B) qRT-PCR for *Nkx2-1* and indicated representative antioxidant gene mRNAs using cDNAs from *Nkx2-1*^fl/fl^ (N=8) and *Nkx2-1*^ΔT^ (N=4) mouse thyroids as described in (A). Data are presented as mean ± SD from individual mouse thyroid samples. For each gene, the mean value of *Nkx2-1*^fl/fl^ mouse samples was set as 1. *p<0.05, **p<0.01, ***p<0.005 by Student’s *t*-tests or Welch’s *t*-test. Significantly up-regulated genes with >2 fold increase in *Nkx2-1*^ΔT^ mouse thyroids are marked with red rectangles. (C) Representative pictures of histology of *Nkx2-1*^fl/fl^ and *Nkx2-1*^ΔT^ mouse thyroids in (A). Note that less number of follicles with irregular shapes are seen in *Nkx2-1*^ΔT^ mouse thyroid as compared to *Nkx2-1*^fl/fl^. H&E stain. X10 objective. Bar, 100 μm. (D) qRT-PCR for gene mRNAs using the cDNAs used in (B). Low, low expression of *Duox1* (See Materials & Methods). For each gene, the mean value of *Nkx2-1*^fl/fl^ mouse samples was set as 1. *p<0.05 by Student’s *t*-tests. (E) qRT-PCR for indicated gene mRNAs using cDNAs from *Nkx2-1*^fl/fl^ mouse thyroids in CD, CD+RAD, IDD, and IDD+RAD groups used in Figure. 2D. Data are presented as mean ± SD from individual mouse thyroid samples (N=3 for each group). Low, low expression of *Duox1* (See Materials & Methods). For each gene, the mean value of CD group thyroids was set as 1. *p<0.05, ***p<0.005 for comparison against CD group thyroids by Dunnett’s multiple comparison tests.

Histological examination revealed that *Nkx2-1*^ΔT^ mouse thyroids in CD group display abnormally compromised architecture of glands with irregular shapes as reported in our previous study [12] (Figure 4C). In addition, focal hyperplasia was observed in some old *Nkx2-1*^ΔT^ mice (Figure S4; Table S1). It should be noted that heterozygous thyroid-specific deletion of *Nkx2-1* resulted in normal thyroid gland architecture and unaltered thyroid adenoma incidence in IDD+RAD group (Figure S5; Table S2), suggesting that the histological anomaly caused by homologous deletion of *Nkx2-1* accounts for the increase in thyroid neoplasm development. Based on this, we thought that the expression of thyroid markers may be altered, possibly as compensation of compromised functions of the abnormal thyroid. In fact, among the thyroid hormone synthesis markers, expression of *Slc5a5* (encoding sodium/iodide symporter, also known as NIS) and *Tpo* (thyroid peroxidase) was found up-regulated in *Nkx2-1*^ΔT^ mice under CD (Figure 4D). Similarly, up-regulation of *Slc5a5* and *Tpo* was induced by iodine deficiency (Figure 4E), suggesting a possibility that disturbance of thyroid functions related to hormone synthesis might be involved in higher ROS production.

### Negative regulation of *Acox2* oxidase expression by NKX2-1 in thyroids

Whether the expression of oxidative stress-related molecules is regulated by NKX2-1 in the thyroid was next examined. From 356 up-regulated genes (>2 fold) in *Nkx2-1*^ΔT^ thyroids, genes with the ontology of “oxidase” were extracted and selected using Database for Annotation, Visualization and Integrated Discovery (DAVID [30, 31]), resulting in the identification of *Acox2* (one subtype of Acyl-coA oxidase, ACOX), *Noxa1* (NADPH Oxidase Activator 1) and *Noxo1* (NADPH Oxidase Organizer 1, Figure. 5A). The latter two are activators of NADPH Oxidase 1 (NOX1) [32], which was not substantially expressed in *Nkx2-1*^ΔT^ thyroids according to the RNA-seq data, therefore these genes were not examined further. ACOX is localized in peroxisome and has three major subtypes (ACOX1, ACOX2, and ACOX3). They are responsible for beta-oxidation of fatty acid and generate H_2_O_2_ as a byproduct [33]. RNA-seq results of *Nkx2-1*^fl/fl^ and *Nkx2-1*^ΔT^ thyroids showed significant up-regulation of only *Acox2* gene among the three subtypes in *Nkx2-1*^ΔT^ thyroids (Figure 5B). Higher *Acox2* expression in *Nkx2-1*^ΔT^ thyroids was consistently observed under various conditions (Figures 5C and S6), suggesting a possibility that NKX2-1 may regulate expression of *Acox2* in thyroid follicular cells.

**Figure 5.**
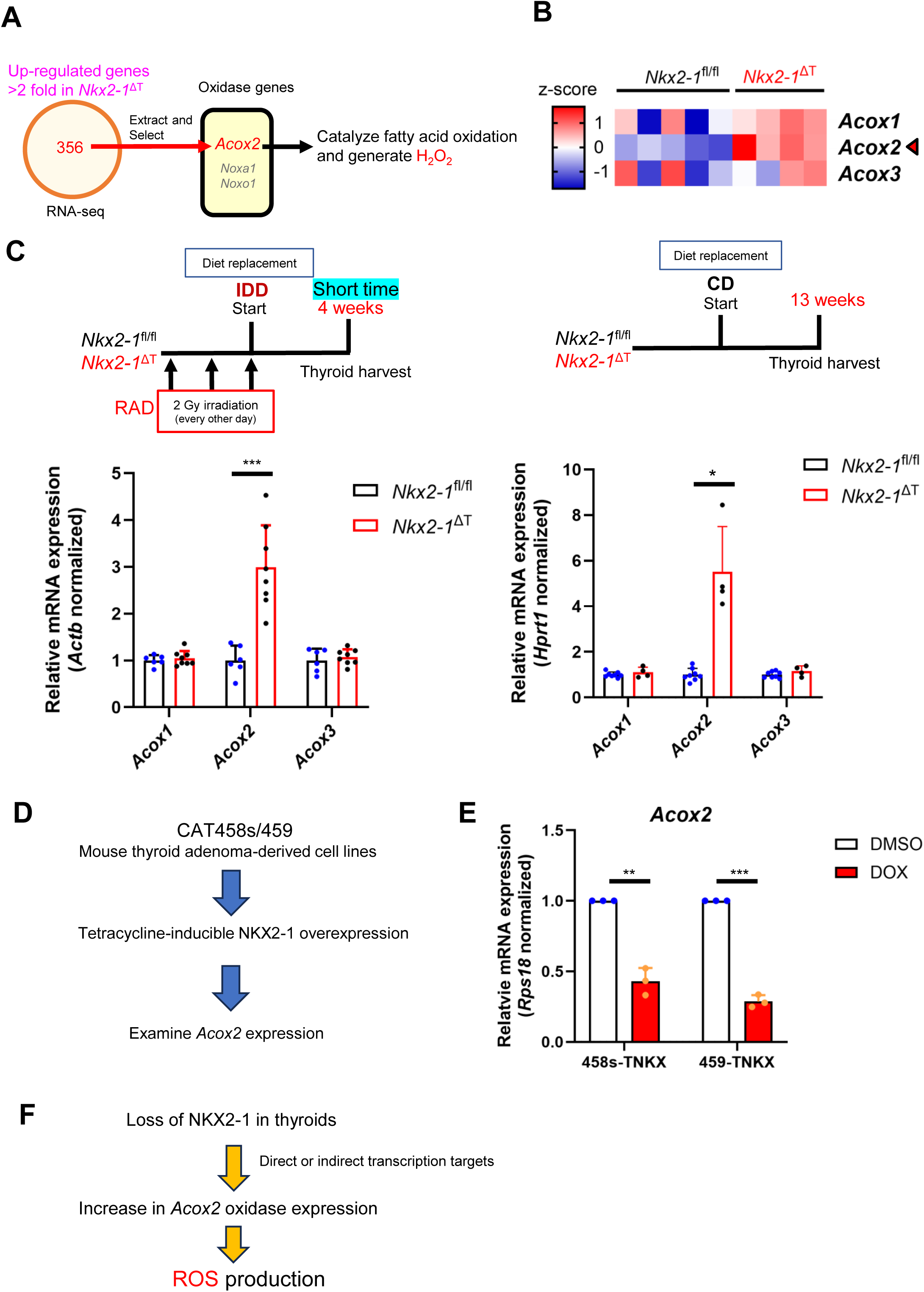
Regulation of *Acox2* oxidase expression by NKX2-1 in mouse thyroids. (A) Schematic representation of identification of *Acox2* in up-regulated oxidase genes in *Nkx2-1*^ΔT^ mouse thyroids using the RNA-seq data as compared with *Nkx2-1*^fl/fl^, as described in Figure 3A. (B) Heatmap of *Acox1*, *Acox2* and *Acox3* using the RNA-seq data in (A). A red arrow indicates significant up-regulation in *Nkx2-1*^ΔT^. (C) qRT-PCR for *Acox1*, *Acox2* and *Acox3* gene mRNAs in mouse thyroids collected at 4 weeks under IDD+RAD (see Figure 3D) and those collected at 13 weeks under CD (see Figure 4B). For each gene, the mean value of *Nkx2-1*^fl/fl^ mouse samples was set as 1. *p<0.05, ***p<0.005 by Welch’s *t*-tests. (D) Schematic representation for the examination of *Acox2* expression regulated by NKX2-1 in mouse thyroid cell lines. (E) qRT-PCR for *Acox2* gene mRNA levels in 458s- and 459-TNKX cells treated with DMSO or DOX for three days. For each sample set, the value of 458s- or 459-TNKX cells with DMSO was set as 1 and the value of 458s- or 459-TNKX cells with DOX was represented as fold change. **p<0.01, ***p<0.005 by paired *t*-tests. (F) Schematic representation of ROS production by up-regulation of *Acox2* induced by loss of NKX2-1 in mouse thyroids.

We previously established cell lines from amitrole-induced mouse thyroid adenomatous nodules [34]. Using these cells, tetracycline-inducible NKX2-1-overexpressing cells, 458s- and 459-TNKX were established [34], which were utilized to examine the regulation of *Acox2* by NKX2-1 (Figure 5D). Suppression of *Acox2* expression by NKX2-1 was observed in both 458s- and 459-TNKX cells (Figures 5E and S7), suggesting that NKX2-1 negatively regulates expression of *Acox2* in mouse thyroid follicular cells. These findings suggest that up-regulation of ACOX2 oxidase by loss of NKX2-1 could also contribute to higher ROS production in *Nkx2-1*^ΔT^ mouse thyroids. The findings are summarized in a proposed model (Figures 5F and Figure 6).

**Figure 6.**
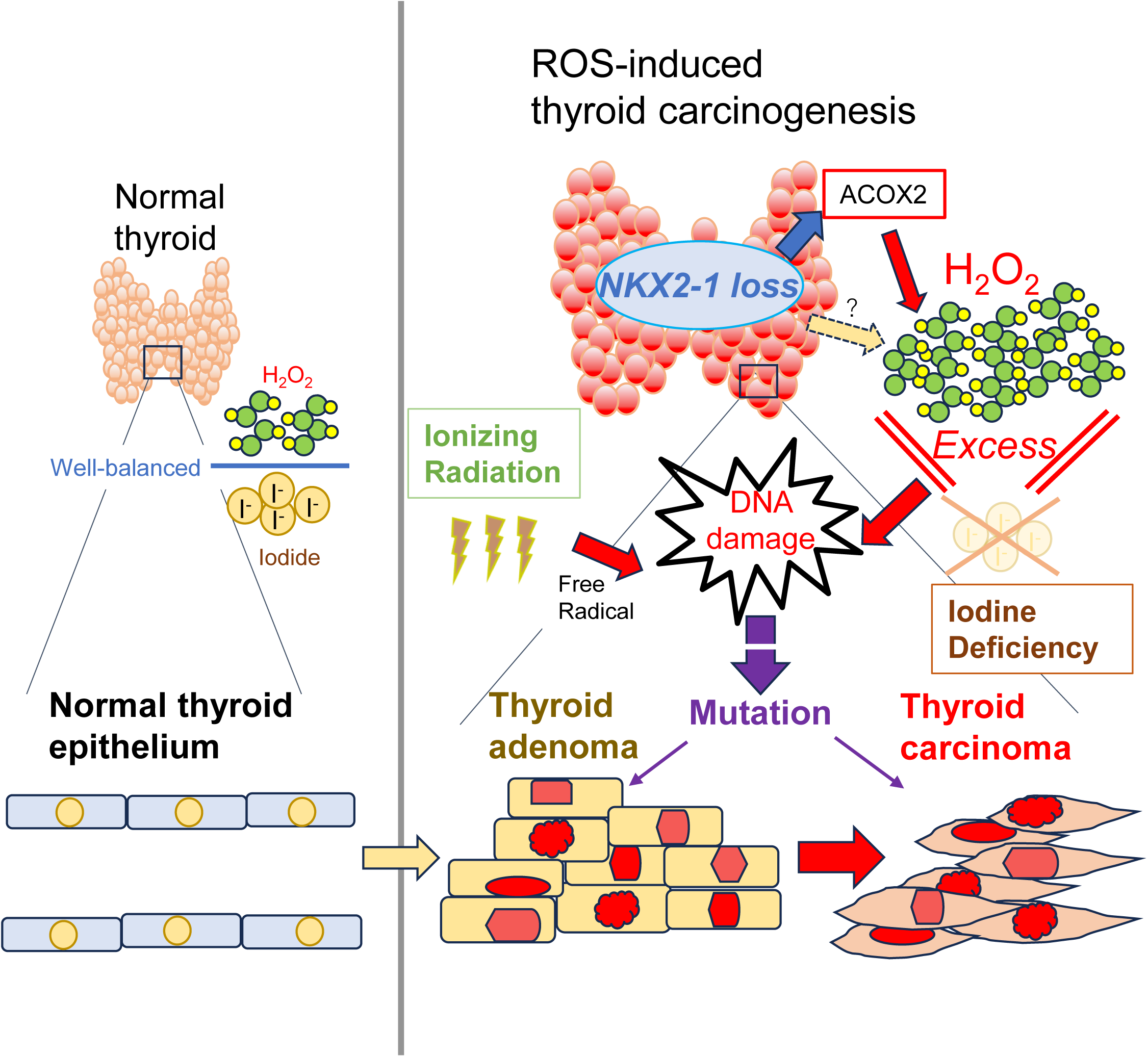
Identification of NKX2-1 as a tumor suppressor in 679 ROS-induced thyroid carcinogenesis. Schematic representation of a proposed model of ROS-induced thyroid carcinogenesis based on our findings. In a normal thyroid, the amounts of H2O2 and iodide are well-balanced and oxidized iodide are used for coupling with thyroglobulin for thyroid hormone synthesis (See References Kohre [1] and Pashcke [20]). Iodine deficiency supposedly leads to imbalance of the iodide oxidation and results in excess H2O2 in follicular cells. Ionizing radiation produces free radicals which eventually causes DNA damage together with H2O2 resulting in DNA mutations involved in development of thyroid adenoma and carcinoma. In addition to these known risk factors for thyroid carcinogenesis, loss of NKX2-1 up-regulates ACOX2 oxidase expression and this may contribute to overproduction of ROS. Loss of NKX2-1 facilitates ROS-induced thyroid carcinogenesis and it was identified as a risk factor for thyroid carcinogenesis.

## Discussion

Amplification of *NKX2-1* gene in lung adenocarcinoma is thought to give advantage on cell survival during carcinogenesis, while tumor suppressive role of NKX2-1 in lung cancer progression has been highlighted [13]. In thyroid cancer, expression of NKX2-1 is generally correlated with the degree of differentiation [35, 36] and is considered as a differentiation marker for diagnosis. Clarification of the role of NKX2-1, the common transcription factor with lung and thyroid cancer, is an imperative need for understanding the similarities and differences in carcinogenesis process between the two different tissues. In this study, the effects of loss of NKX2-1 on thyroid carcinogenesis induced by two known thyroid-specific risk factors, iodine deficiency and ionizing radiation were examined that were found to induce oxidative stress in an additive fashion. Adenoma and carcinoma development was promoted by loss of NKX2-1 in our model (Tables 1 and S2). It should be noted however, that all carcinomas developed in our study were adenocarcinomas in a non-papillary pattern some of which supposedly arise from adenomas as proposed in multi-step carcinogenesis [7]. Classic PTC development is thought to directly arise from normal follicular cells [7], and thus the use of different study models may be required to understand the effects of NKX2-1 on classic PTC development.

SODs work specifically against superoxide anion (O ^・-^) among a variety of free radicals and have three subtypes (SOD1, SOD2 and SOD3) in mammals [21]. They have distinct metal ion co-factors and localizations, that is, Cu/Zn SOD1 in cytoplasm, Mn SOD2 in mitochondria matrix, and Cu/Zn SOD3 in extracellular space [21]. Their actions are thought to be strictly regulated in a spatial manner. Generated H_2_O_2_ through a reaction of O_2_ ^・- with SODs further needs to be detoxicated by direct-acting antioxidant enzymes, i.e. CAT, GPXs and PRDXs, to prevent DNA damage [21, 37, 38]. CAT does not have any subtypes in mammals and is mainly localized in peroxisome in which beta-oxidation of fatty acid takes place, while GPXs have eight subtypes (GPX1-8) and PRDXs have six subtypes (PRDX1-6); they are more ubiquitously distributed depending on the subtypes with their specific organs and extra- or intracellular spaces^ [25–27]. We identified SOD3 and GPX2 as the key up-regulated antioxidants in response to oxidative stress in the thyroid in our study model. It is postulated that surplus H_2_O_2_ supposedly caused from iodine deficiency induced up-regulation of SOD3 as a signaling molecule, as demonstrated by the previous report [22]. However, up-regulation of *Gpx2* by iodine deficiency was not observed in Balb/c mice in the same study [22]. Taking it into consideration that GPX2 is abundant in the gastrointestinal tract and targets mainly lipid hydroperoxide [26], mouse strain and condition-specific induction of antioxidants might be present.

Besides SOD3 and GPX2, other direct-acting antioxidant enzymes, SOD1, SOD2, CAT, and PRDX6 were found up-regulated at protein level in IDD and IDD+RAD groups in spite of unchanged or decreased mRNA expression at 13 weeks after diet replacement (Figures 2D and S8). On the contrary, at an earlier time point of 4 weeks after diet replacement, SOD3 was already clearly induced by iodine deficiency even though up-regulation of other proteins, SOD1, CAT, and PRDX6 was not observed (Figure S9C). Up-regulation of CAT protein was also detected in IDD+RAD-treated *Nkx2-1*^ΔT^ thyroids under various conditions (Figures 3I and S3B), suggesting that CAT can also serve as a representative oxidative stress marker in the thyroid based on its essential function as an anti-oxidative stress defense agent [39]. These results suggest a possibility for posttranscriptional regulation of some antioxidant proteins under IDD and/or IDD+RAD.

ACOX has three subtypes with distinct distributions and target lipids. ACOX1 is in charge of very long straight-chain fatty acids, while ACOX2 and ACOX3 have specificities to branched chain fatty acids [33]. ACOX2 was reported to be specifically involved in bile acid synthesis [40]. Consistent with this, expression of *Acox2* was detectable but not abundant in mouse thyroid according to the RNA-seq data (RSEM for *Nkx2-1*^fl/fl^ mice <100). Considering consistent up-regulation of *Acox2* in *Nkx2-1*^ΔT^ thyroids under various conditions (Figures 5C and S6), it is speculated that the loss of NKX2-1 induces ectopic expression of ACOX2 in the thyroid, which might contribute to dysregulation of thyroid function and homeostasis. On the other hand, iodine deficiency for 13 weeks did not significantly up-regulate *Acox2* expression in *Nkx2-1*^fl/fl^ mouse thyroids (Figure S10), suggesting that the loss of NKX2-1, but not iodine deficiency-derived thyroid dysfunction, affects *Acox2* expression. This is consistent with the fact that *Acox2* expression is suppressed by NKX2-1 overexpression in mouse thyroid adenoma-derived cells (Figures 5E and S7). Further studies are required to understand the role of ACOX2 in thyroid function and how its expression is regulated by NKX2-1.

Thyrotropin (TSH) is reported to up-regulate expression of thyroid markers [41] and stimulate production of H_2_O_2_ in thyroid follicular cells [42–45]. Iodine deficiency is known to cause elevation of TSH as a negative feedback [17]. Therefore, it could be possible that TSH is involved in higher ROS production in *Nkx2-1*^fl/fl^ mouse thyroids in IDD group with the finding of up-regulation of *Tshr*, *Slc5a5* and *Tpo* (Figure 4E). However, we previously reported only a small portion of *Nkx2-1*^ΔT^ mice had clearly elevated TSH [12]. Further, similar up-regulation of *Tshr*, *Slc5a5* and *Tpo* in *Nkx2-1*^ΔT^ thyroids was not consistently observed among different schedules of diet replacement and radiation (Figures 4D and S11). This suggests that unknown other factors than TSH might play a role in up-regulation of these markers and possible subsequent ROS overproduction in *Nkx2-1*^ΔT^ thyroids. Only around 50% reduction of NKX2-1 expression at maximum was achievable using this model as reported [12], thus different designs of mouse models would be required to further identify targets of NKX2-1 in ROS production and fully clarify their functions.

In conclusion, we found a critical role of NKX2-1 in ROS-mediated thyroid carcinogenesis. Our findings would give a further understanding of thyroid-specific carcinogenesis, especially multi-step thyroid carcinogenesis process with adenoma-carcinoma sequence.

## Materials & Methods

### Mice with gamma-ray irradiation and diet replacement

Mice used in this study harbored either one or two *Nkx2-1*-floxed allele(s) with or without *TPO*-Cre, namely *Nkx2-1*^fl/wt^, *Nkx2-1*^fl/fl^, *Nkx2-1*^fl/wt^;*TPO*-Cre (*Nkx2-1*^HΔT^), and *Nkx2-1*^fl/fl^;*TPO*-Cre (*Nkx2-1*^ΔT^). Mice were backcrossed to C57BL/6 (Chrales River Laboratories, MA, USA) for 6 (for Figures 1D, 1E and S4; Tables 1 and S1, and all the qRT-PCR, RNA-seq, Western blotting results shown in Figures 2-5, S2, S3, S6, and S8-S11) or 7 (for Figure 4C and S5; Table S2) times. *Nkx2-1*^fl/fl^ and *Nkx2-1*^ΔT^ mice were used for the experiments as control and thyroid-specific *Nkx2-1* conditional knockout, respectively, unless otherwise mentioned. Note that *Nkx2-1*^ΔT^ mouse thyroids are hypomorphic with various degrees of deletion of *Nkx2-1* depending on each cell, making an average of ∼50% deletion and cells that lost NKX2-1 expression degenerate as previously described [12, 15]. All animal studies were performed under the protocols approved by the National Center Institute Animal Care and Use Committee (LM-024 and 077).

### Experiments for radiation and iodine deficiency were carried out as follows

1. Radiation (RAD): Total body irradiation with gamma rays (2 Gy (195-200 cGy)) was performed using Gammacell 40 Exactor (Best Theratronics Ltd., Ontario, Canada) a total of three times every other day.
2. Diet replacement: Mouse diet was changed right after the last total body irradiation on the same day to Control diet (CD; Rodent Diet, AIN-93G 1/2" Pellets, Bio-Serv, NJ, USA) or Iodine deficient diet (IDD; Rodent Diet, AIN-93G, Modified, Iodine Deficient, 1/2" Pellets, Bio-Serv).

Four experimental groups were used as follows:

1. CD group; Control diet without Radiation (RAD), 2) CD+RAD group; Control diet with Radiation, 3) IDD group; Iodine deficient diet without Radiation, and 4) IDD+RAD group; Iodine deficient diet with Radiation (Figure. 1B). Male mice of various genotypes were randomly assigned to a group described above and the diet was changed at the age of 5.0 to 8.0 weeks old right after RAD on the last day or without it. Mice were euthanized over a year (1.08 to 1.28 years for Table S2 and 1.10 to 1.30 years for the rest) after the start of special diet for the histological analysis for thyroid lesion incidence.

### Cell lines and culture

CAT458s and 459 cells were established from male mouse thyroid adenomatous nodules in our laboratory as previously reported [34]. These cells were used to obtain stable cell lines, named 458s- and 459-TNKX with tetracycline-inducible expression of rat *Nkx2-1* 458s- and 459-TNKX with tetracycline-inducible expression of rat *Nkx2-1* [34]. They were cultured in DMEM/F12 medium (Corning, NY, USA) containing 10% fetal bovine serum (FBS, GeminiBio, CA, USA) and Antibiotic:Antimycotic Solution (Gibco/Thermo Fisher Scientific, MA, USA). Cells were maintained and grown in a 5% CO_2_ atmosphere at 37°C. Whole genome sequencing (WGS) of the parental CAT458s and 459 was performed as previously reported [34]. The WGS data can be considered as an authentication of these cell lines. Doxycycline (Merck Millipore, MA, USA) or DMSO (R&D systems, MN, USA) was added to cells seeded in a 6 well-plate to examine the effect of NKX2-1 overexpression.

### RNA-seq

Mouse thyroid tissues were immersed in RNA*later* solution (Thermo Fisher Scientific) immediately after resection, and homogenized using Precellys 24 tissue homogenizer (Bertin Technologies, Montigny-le-Bretonneux, France). Total RNAs were extracted using QIAshredder (QIAGEN, Hilden, Germany), RNeasy Plus mini kit (QIAGEN) and RNase-Free DNase Set (QIAGEN). Samples with RIN >7.0 measured with TapeStation were subjected to RNA-seq. Nine total RNA-Seq samples (five samples from *Nkx2-1*^fl/fl^ thyroids and four samples from *Nkx2-1*^ΔT^ thyroids) were pooled and sequenced on NovaSeq Xplus 10B using Illumina Stranded Total RNA Prep, Ligation with Ribo-Zero Plus and paired-end sequencing (Illumina, San Diego, CA). RNA-seq generated 477 to 610 million high-quality reads per sample (>91% above Q30). After trimming with Cutadapt, reads were mapped to the reference mouse genome (mm10) using STAR (v2.7.11b). Mapping metrics were calculated with Picard, yielding an average 82% mapping rate, >49% unique alignment, and 5.2% ribosomal bases. Coding, UTR, and total mRNA bases represented 22–34%, 36–56%, and 70–80% of reads, respectively. Library complexity (Picard MarkDuplicates) showed 6–27% non-duplicate fragments. Gene expression was quantified using the STAR/RSEM pipeline to generate TPM and FPKM values. Differential expression analysis was performed with DEseq2 (over 500 total RSEM gene counts for each group). The RNA-seq data has been deposited in the NCBI Gene Expression Omnibus (GEO) and can be accessible through GEO Series accession number GSE310837.

### Gene Set Enrichment Analysis (GSEA)

Gene Set Enrichment Analysis (GSEA) of RNA-seq data used the gene sets from the Mouse Molecular Signatures Database (MSigDB) [28]. Enrichment results from WikiPathways (a subset of Canonical pathways) and mouse-ortholog hallmark gene sets (MH) [46] are shown in Figure. 3B; Tables S3 and S4 (FDR<0.25). Heatmaps were generated using z-scores from log2-transformed RSEM gene counts with GraphPad Prism (GraphPad Software, MA, USA).

### Ontology knowledgebase analysis

Gene ontology was analyzed using Database for Annotation, Visualization and Integrated Discovery (DAVID) [30, 31]. Genes annotated as “Oxidase” by GO ontology and KEGG Pathway ontology were extracted from the up-regulated gene list in *Nkx2-1*^ΔT^ mouse thyroids (>2 fold), and the genes which could directly contribute to oxidative stress were selected (Figure 5A).

### Quantitative reverse transcription PCR analysis (qRT-PCR)

Total RNAs from mouse thyroids, and 458s- and 459-TNKX cells were prepared using QIAshredder (QIAGEN) and RNeasy Plus mini kit (QIAGEN). Extracted RNAs used for RNA-seq were also subjected to the analysis. cDNA was synthesized using SuperScript IV First-Strand Synthesis System (Thermo Fisher Scientific) using 68.6 ng to 1 μg RNAs obtained from mouse thyroids (the same amount was used for the same sample set) and 1 μg from 458s/459-TNKX cells. qRT-PCR was run as previously reported using PerfeCTa SYBR Green FastMix, Low ROX (Quantabio, MA, USA) and QuantStudio 7 Flex Real-Time PCR System using standard curve method (Thermo Fisher Scientific) [34]. For tissues, multiple mouse thyroids were used per group (N>=3), while cells were collected at multiple independent occasions (N=3, Figure 5E) and biological replicates were used per one sample (N=3, Figure S7). Values for gene expression per sample were calculated using the average qRT-PCR values obtained from two or three wells, and were normalized to *Hprt1*, *Actb* or *Rps18* mRNA. Primers used in this study are listed in Table S5.

### Immunoblotting

Thyroid lysates from *Nkx2-1*^fl/fl^ and *Nkx2-1*^ΔT^ mice were prepared in RIPA lysis buffer (Merck Millipore) containing cOmplete Mini EDTA-free Protease Inhibitor Cocktail (Roche, Basel, Switzerland) with PhosStop (Roche). Westen blotting was performed as previously reported using Criterion TGX Precast Gels and the Turbo Trans-Blot Transfer System (BIO-RAD, CA, USA) [34]. TBS buffer containing BSA fraction V (Merck Millipore) in addition to Tween-20 was used for blocking. H_2_O_2_ and Restore Western Blot Stripping Buffer (Thermo Fisher Scientific) were used for the inactivation of HRP signal and stripping of the membrane, respectively. ChemiDoc Imaging System with Image Lab software (BIO-RAD) was used for visualization, capture of images and quantification of the obtained bands with a gamma value 1.15 of the original Auto Scale settings. Pierce BCA Protein Assay Kit (Thermo Fisher Scientific) was used for protein concentration measurement. Multiple individual mouse thyroid samples (N>=3) were used for comparison. Antibodies used in this study are listed in Table S6.-

### Histological analysis

Mouse thyroid tissues were fixed in 10% buffered formalin and embedded in paraffin, and sections were stained with hematoxylin and eosin (H&E), performed by Histoserv, Inc (MD, USA) or in our laboratory at the National Cancer Institute (NCI). A board certified veterinary pathologist (JMW) examined the thyroid sections and classified thyroid lesions as noted in the Results based on the standard international veterinary and toxicology nomenclature [47].

### TCGA PanCancer Atlas dataset and statistical analysis

Thyroid Carcinoma (TCGA, PanCancer Atlas) dataset was utilized through cBio Cancer Genomic Portal (cBioPortal, https://www.cbioportal.org/) [48, 49] for prognosis analysis. Among 499 patients, 7 patients annotated as “Other” tumor type were removed from the analysis and only thyroid papillary carcinoma patients (Figure S1). mRNA expression data was not available for 2 patients, and 6 patients deceased because of events not related to the cancer. Relationship of disease-specific survival and *NKX2-1* expression in the primary lesion was analyzed in 484 PTC patients. *NKX2-1* expression was classified using z-scores relative to normal samples as follows: z-score>1; High, 1=>z-score>-1; Intermediate, −1=>z-score; Low. Tumor type graph, disease-specific survival graph and p-value by Logrank test were generated by cBioPortal (Figure 1A).

### Statistical analysis

Two-tailed Student’s *t*-tests, paired *t*-tests or Welch’s *t*-tests were used for comparison of two samples. Dunnett’s and Tukey’s multiple comparison tests were applied for comparison of multiple groups against control group and comparison of every group, respectively. Fisher’s exact tests were applied regarding the comparison between two-group (*Nkx2-1*^fl/fl^ and *Nkx2-1*^ΔT^) in histology analysis (Tables 1 and S1). Chi-square test for multiple groups was used for comparison among three groups (*Nkx2-1*^fl/wt^ or *Nkx2-1*^fl/fl^, *Nkx2-1*^HΔT^and *Nkx2-1*^ΔT^, Table S2). p-values for RNA-seq analyses were obtained using the statistical modeling framework implemented in DESeq2, which applies the Wald test for differential gene expression comparisons. Results were considered statistically significant at p<0.05.

## Acknowledgments

RNA-seq was conducted at the Sequencing Facility at NCI, National Institutes of Health (NIH). Mice under Control diet and Iodine deficient diet were maintained at the animal facility run by the NIH Veterinary Resources Program. We would like to thank Jorge A. Paiz and Nabanita Kundu of Cancer Innovation Laboratory of NIH for genotyping and maintenance of mice. The results for the thyroid cancer patients shown here are in whole or part based upon data generated by the TCGA Research Network: https://www.cancer.gov/tcga. Part of this work was presented at the American

Association for Cancer Research (AACR) Annual Academic Meeting 2021 (https://doi.org/10.1158/1538-7445.AM2021-2661). This research was supported in whole by the Cancer Innovation Laboratory, Center for Cancer Research, National Cancer Institute, National Institutes of Health Intramural Research Program (SK, project number ZIA BC 005522) and federal funds from the National Cancer Institute, National Institutes of Health, under contract HHSN261200800001E. The contributions of the NIH author(s) were made as part of their official duties as NIH federal employees, are in compliance with agency policy requirements, and are considered Works of the United States Government. However, the findings and conclusions presented in this paper are those of the author(s) and do not necessarily reflect the views of the NIH or the U.S. Department of Health and Human Services.

## CRediT Author Contributions

Yo-Taro Shirai: Conceptualization, Data Curation, Formal analysis, Investigation, Visualization, Writing - original draft, Writing - review and editing. Jerrold M. Ward; Data Curation, Formal analysis, Writing – review and editing. Yoshinori Takizawa: Data Curation, Investigation. Huaitian Liu: Data Curation, Formal analysis, Writing – review and editing. Masaaki Miyakoshi: Investigation, Methodology. Manabu Iwadate: Data Curation, Investigation. Tsubasa Murata: Investigation. Suguru Hayase: Investigation. Shigetoshi Yokoyama: Investigation. Shogo Ehata: Supervision, Writing – review and editing. Shioko Kimura: Conceptualization, Methodology, Funding acquisition, Supervision, Writing – review and editing.

