## Supplemental Information for "Loss of NKX2-1 predisposes thyroid to neoplasm development through regulation of oxidative stress"

**This PDF file includes:**

Figures S1 to S11
Tables S1 to S6
SI References

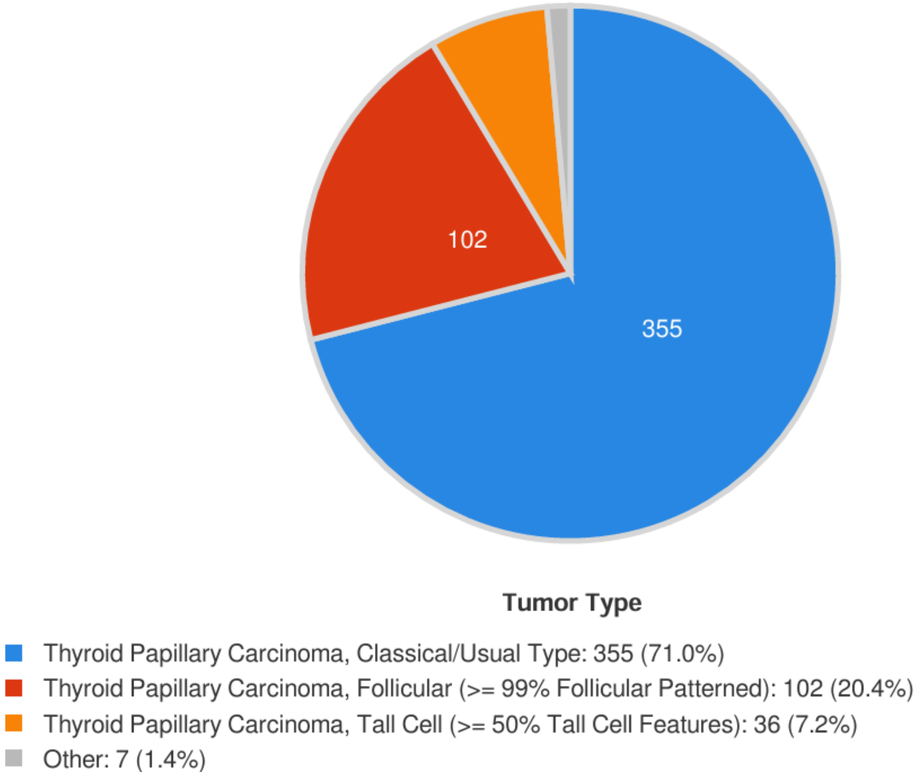


**Figure S1. Tumor types of thyroid cancer in the TCGA dataset of Thyroid Carcinoma PanCancer Atlas.** Details of 500 samples in total (499 primary samples and 1 metastasis sample) of tumor types in the TCGA dataset of Thyroid Carcinoma PanCancer Atlas. One primary sample and one metastasis sample were derived from the same thyroid papillary carcinoma, classical/usual type, patient. 484 primary samples and patients were analyzed for disease-specific survival in Figure 1A after removal of 7 patients classified as “Other” tumor type, 2 patients without the mRNA expression data, and 6 deceased patients due to the reasons not related to the cancer.

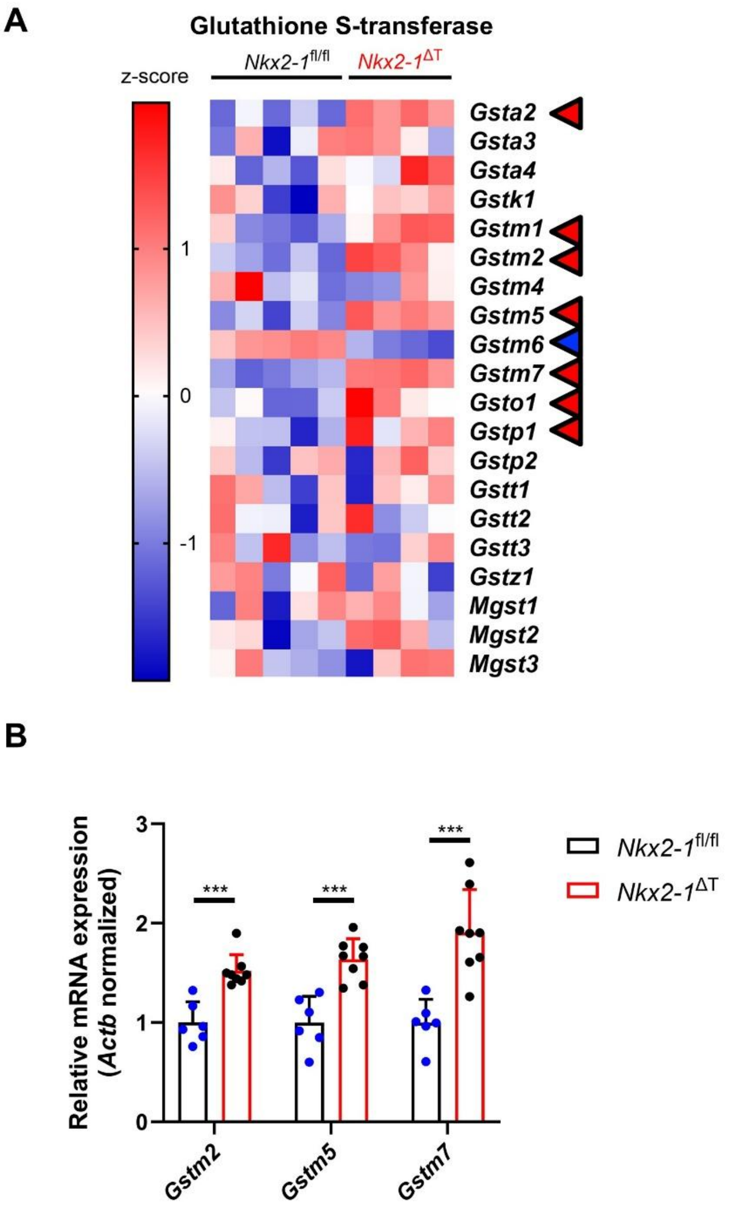


**Figure S2. Up-regulation of Glutathione S-transferase gene expression in *Nkx2-1*$^{\Delta T}$ mouse thyroids with combined exposure to iodine deficiency and radiation 4 weeks after diet replacement.** (A) Heatmap of Glutathione S-transferase genes using the same RNA-seq data as those in Figure 3C. Significantly up-regulated genes or down-regulated gene in *Nkx2-1*$^{\Delta T}$ mouse thyroids are indicated with red arrows or a blue arrow, respectively. (B) qRT-PCR for indicated representative Glutathione S-transferase gene mRNAs using cDNAs from *Nkx2-1*$^{fl/fl}$ (N=6) and *Nkx2-1*$^{\Delta T}$ (N=8) mouse thyroids used in Figure 3D. Data are presented as mean ± SD obtained from individual mouse thyroid samples. For each gene, the mean value of *Nkx2-1*$^{fl/fl}$ mouse samples was set as 1. ***p<0.005 by Student's *t*-tests.

**Figure S3.** *Shirai* et al.

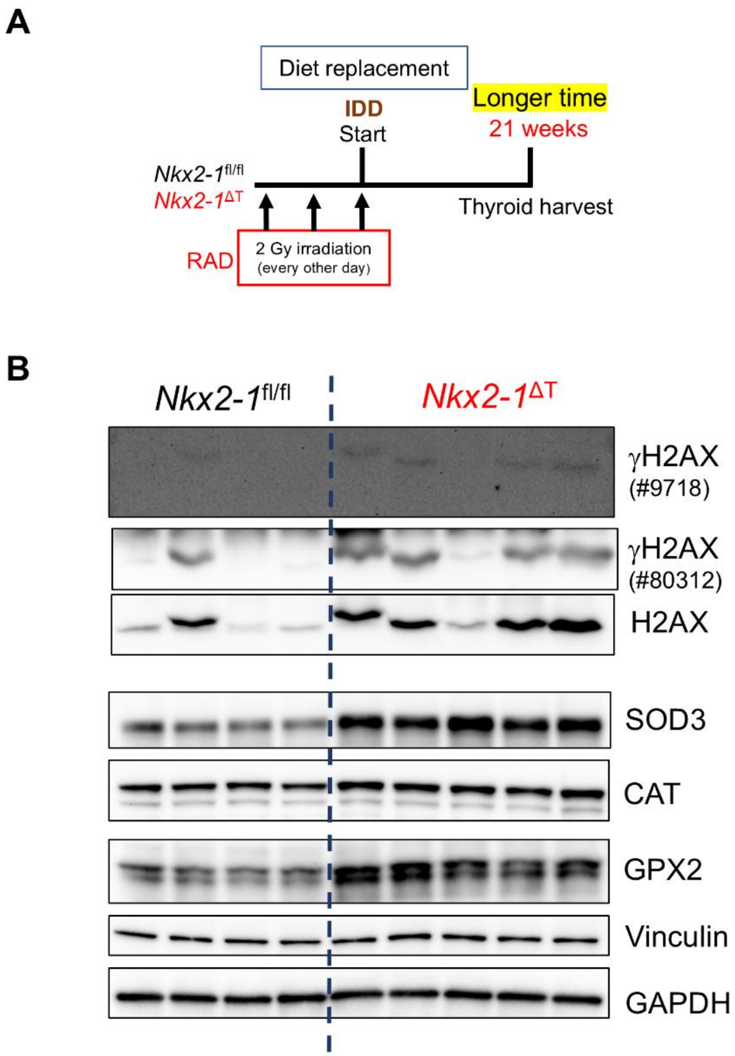


**Figure S3. Enhanced DNA damage and increased expression of antioxidants in IDD+RAD-treated *Nkx2-1*$^{\Delta T}$ mouse thyroids persist for a longer period.** (A) Scheme of time course of *Nkx2-1*$^{fl/fl}$ and *Nkx2-1*$^{\Delta T}$ mouse thyroid collection in IDD+RAD group at 21 weeks after diet replacement. (B) Lysates from *Nkx2-1*$^{fl/fl}$ and *Nkx2-1*$^{\Delta T}$ mouse thyroids in (A) were subjected to Western blotting with the indicated antibodies (CAT, #14097). Blotting images with two different antibodies against γH2AX are shown. Vinculin and GAPDH were used as loading controls.

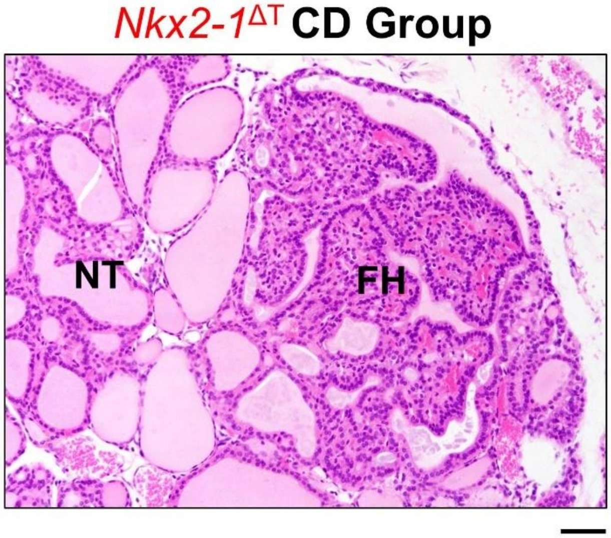


**Figure S4. Histology of focal hyperplasia developed in *Nkx2-1*$^{\Delta T}$ mouse thyroids examined over 1 year after diet replacement.** Representative picture of focal hyperplasia observed in a *Nkx2-1*$^{\Delta T}$ mouse thyroid under Control diet. X20 objective. Hematoxylin and Eosin stain. NT, normal thyroid; FH, focal hyperplasia. Bar, 50 µm.

*Nkx2-1*$^{H\Delta T}$

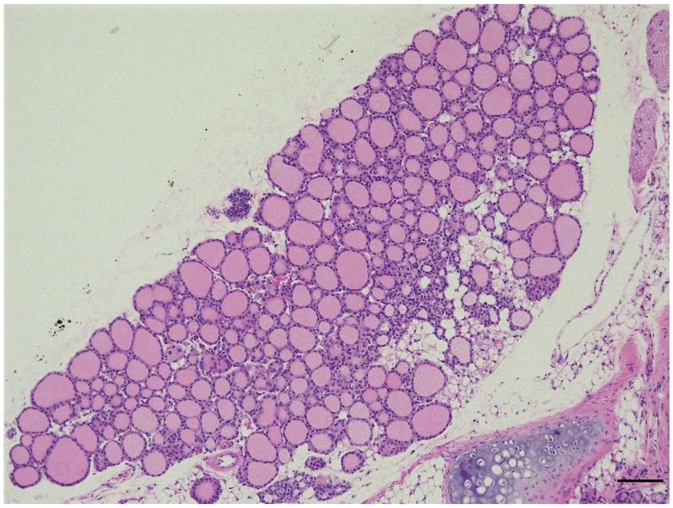

**Figure S5. Histology of thyroid-specific heterozygous deletion of *Nkx2-1* under Control diet.** Representative picture of *Nkx2-1*$^{fl/wt}$;*TPO*-Cre (*Nkx2-1*$^{H\Delta T}$) mouse thyroid 13 weeks after diet replacement with Control diet. X10 objective. Hematoxylin and Eosin stain. Bar, 100 μm.

### Figure S6.

*Shirai* et al.

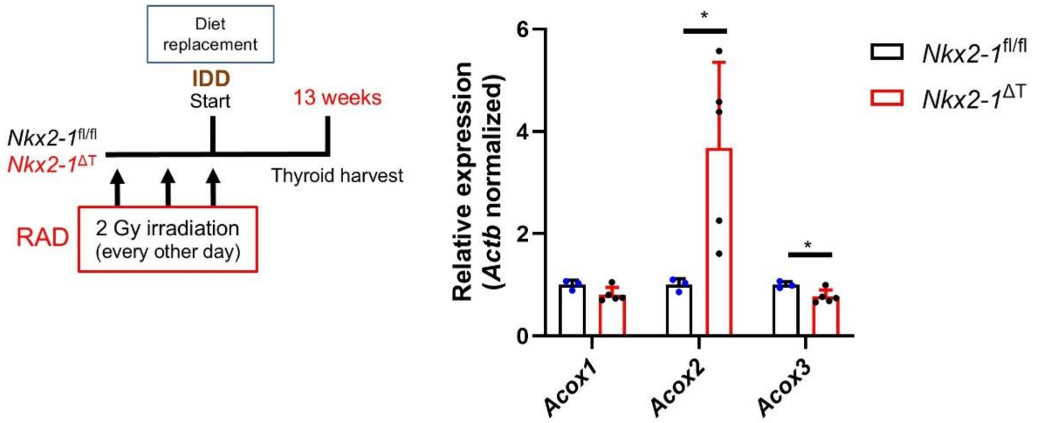


**Figure S6. Up-regulation of *Acox2* in *Nkx2-1*$^{\Delta T}$ mouse thyroids with combined exposure to iodine deficiency and radiation 13 weeks after diet replacement.** Schematic representation of the time course (Left) and qRT-PCR results for *Acox1*, *Acox2* and *Acox3* mRNAs using the cDNAs from *Nkx2-1*$^{fl/fl}$ and *Nkx2-1*$^{\Delta T}$ thyroids used in Figure 3H (Right). Data are presented as mean ± SD from individual mouse thyroid samples. For each gene, the mean value of *Nkx2-1*$^{fl/fl}$ mouse samples was set as 1. *p<0.05 by Student's *t*-test or Welch's *t*-test.

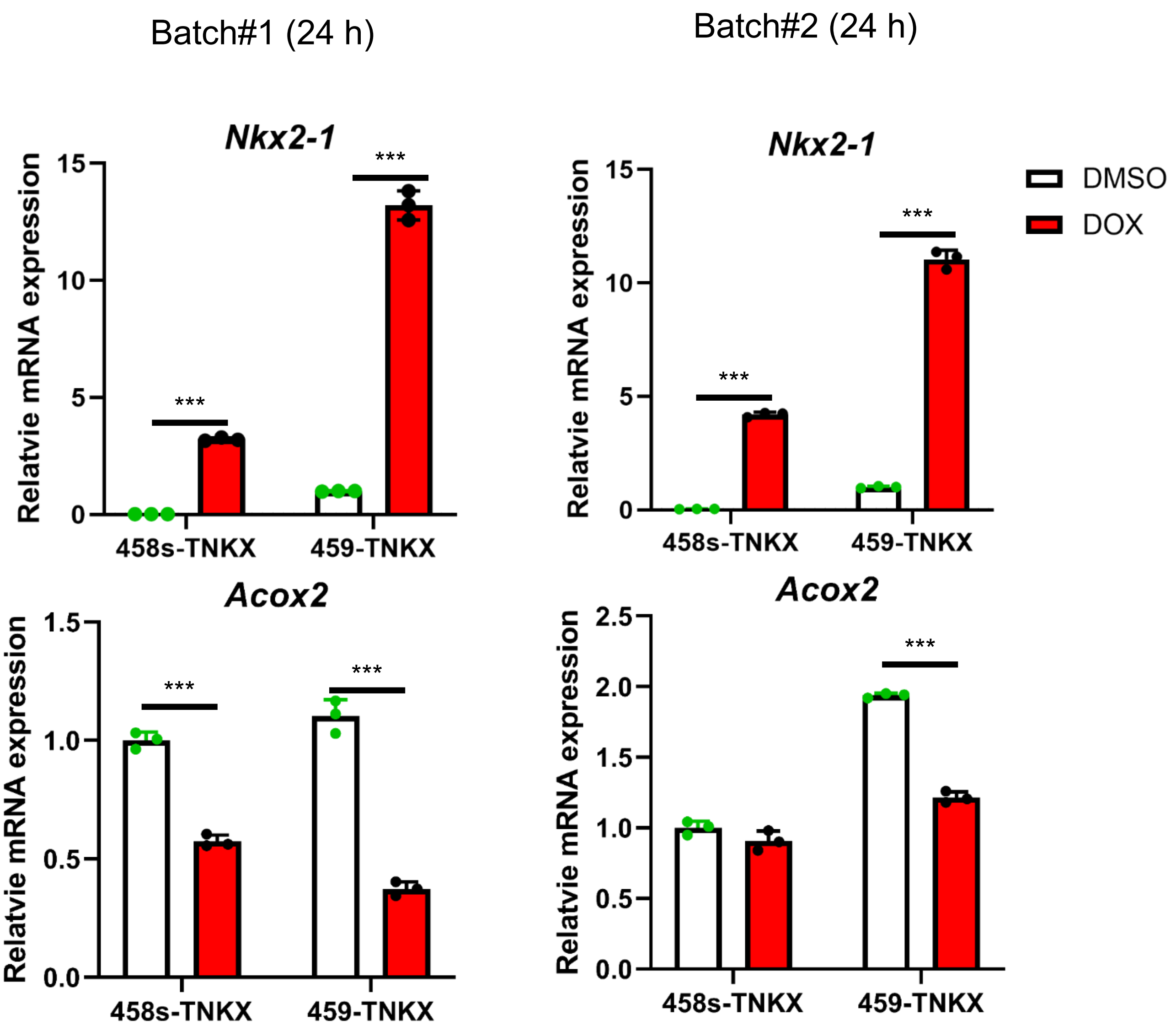


**Figure S7. Changes of *Acox2* expression levels 24 hours after induction of NKX2-1 expression by DOX in 458s/459-TNKX cells.** qRT-PCR for *Nkx2-1* (for mouse/rat) and *Acox2* mRNAs using cDNAs from 458s- and 459-TNKX cells 24 hours after DMSO or DOX treatment. Data are presented as mean ± SD obtained from biological triplicates. Mean value of 458s-TNKX cells treated with DMSO was set as 1. *Rps18* mRNA was used for normalization. ***$p<0.005$ by Student's *t*-tests or Welch's *t*-tests.

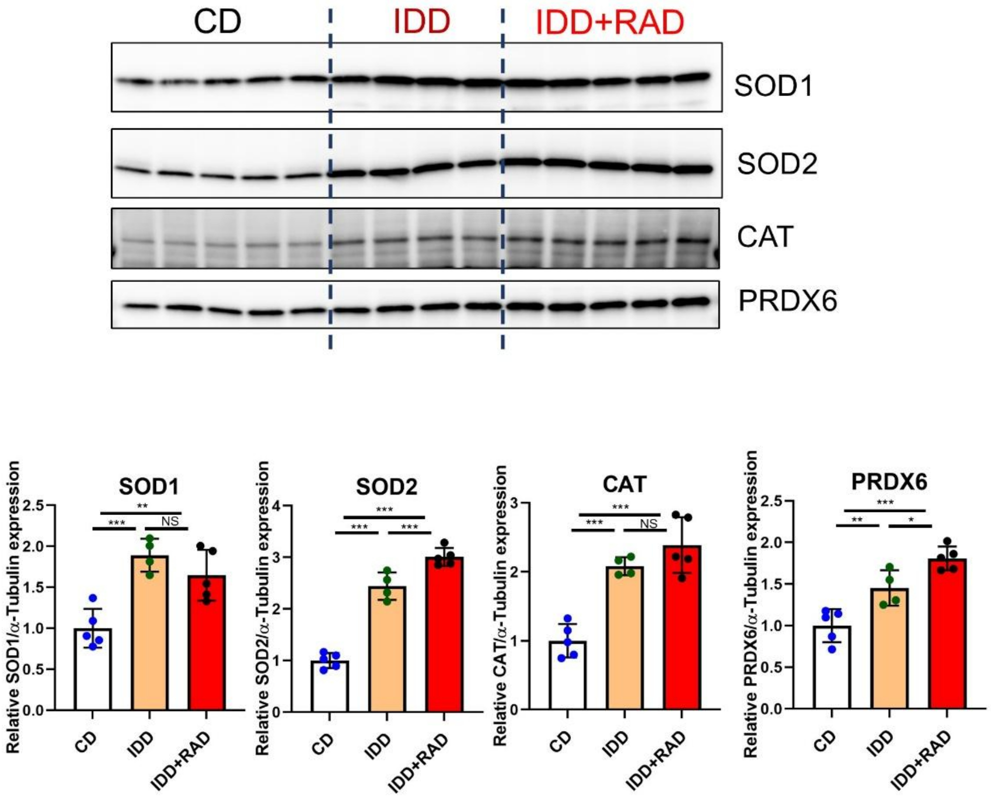


**Figure S8. Changes in expression levels of SOD1, SOD2, CAT, and PRDX6 protein with exposure to iodine deficiency and radiation in *Nkx2-1*$^{fl/fl}$ mouse thyroids at 13 weeks after diet replacement.** Immunoblotting of *Nkx2-1*$^{fl/fl}$ mouse thyroid lysates (used in Figures 2C and 2E) in CD, IDD, and IDD+RAD groups with antibodies against SOD1, SOD2, CAT (ab52477) and PRDX6 (Upper). See β-Actin and α-Tubulin in Figures 2C and 2E as loading controls. Quantification of SOD1, SOD2, CAT or PRDX6/α-Tubulin (Lower). Data are presented as mean ± SD from individual mouse thyroid samples. Mean value of CD group samples was set as 1. *p<0.05, **p<0.01, ***p<0.005 by Tukey's multiple comparison tests. NS, not significant.

#### Figure S9.

*Shirai* et al.

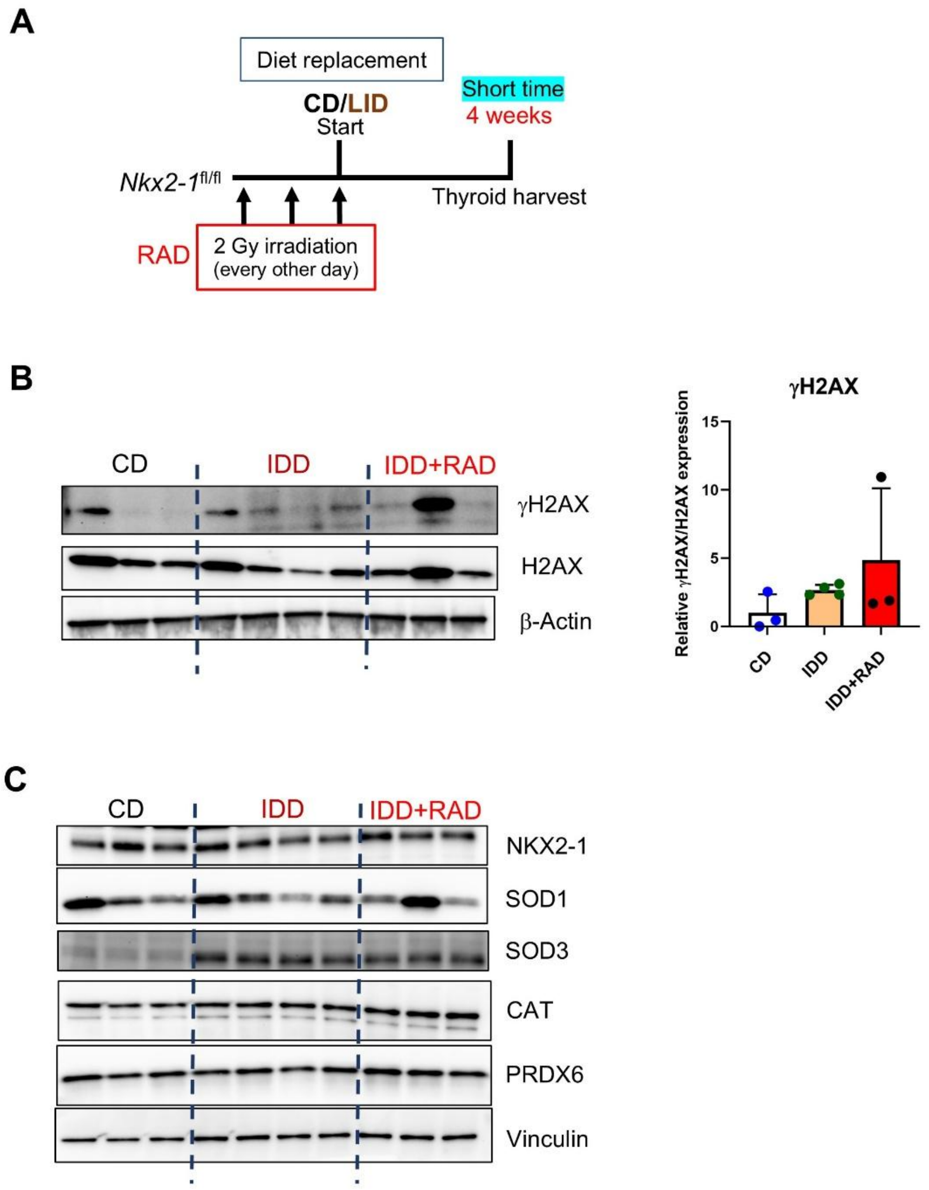


**Figure S9. Up-regulation of SOD3 expression in *Nkx2-1*$^{fl/fl}$ mouse thyroid by iodine deficiency for 4 weeks.** (A) Schematic representation of time course of molecular profile analysis using *Nkx2-1*$^{fl/fl}$ mouse thyroids in CD, IDD, and IDD+RAD groups at 4 weeks after diet replacement. (B and C) Immunoblotting of mouse thyroid lysates in the groups described in (A) with indicated antibodies (γH2AX, #9718; CAT, ab52477). β-Actin and Vinculin were used as loading controls. Quantification of γH2AX/H2AX using Image Lab Software (Right, B). Data are presented as mean ± SD from individual mouse thyroid samples (N=3 or 4). Mean value of CD group samples was set as 1.

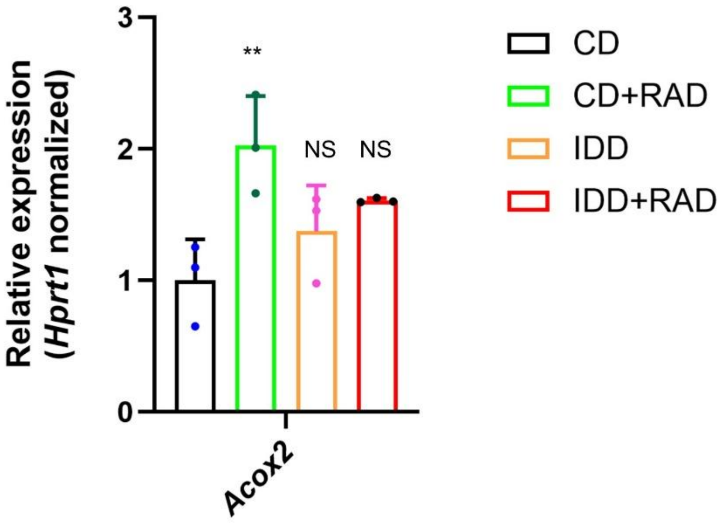


**Figure S10. No significantly altered expression of *Acox2* by iodine deficiency for 13 weeks in *Nkx2-1*$^{fl/fl}$ mouse thyroids.** qRT-PCR for *Acox2* mRNA using cDNAs from *Nkx2-1*$^{fl/fl}$ mouse thyroids in CD, CD+RAD, IDD, and IDD+RAD groups used in Figure 2D. Data are presented as mean ± SD from individual mouse thyroid samples (N=3 for each group). Mean value of CD group thyroids was set as 1. **p<0.01 for comparison against CD group thyroids by Dunnett's multiple comparison tests. NS, not significant.

**Figure S11.** *Shirai* et al.

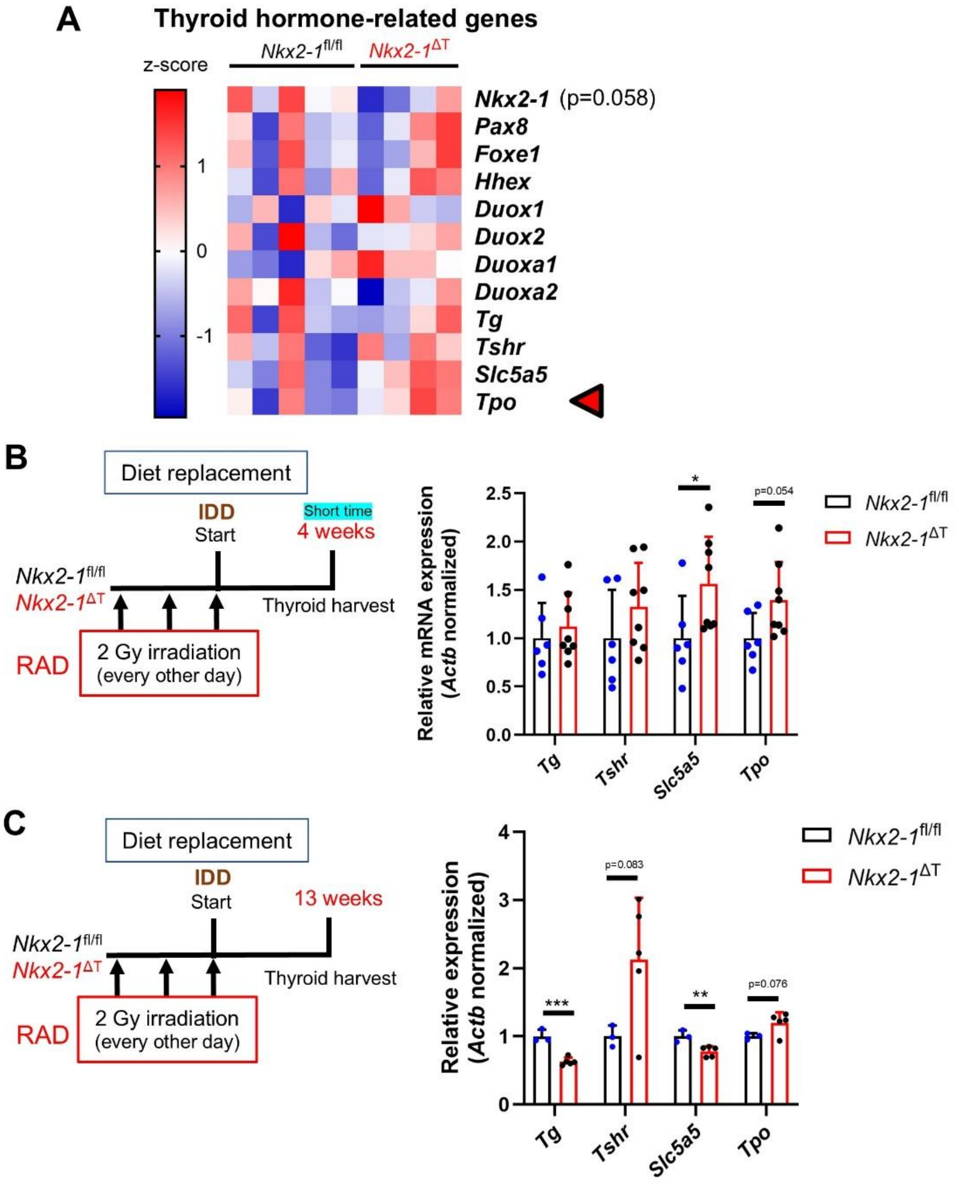


**Figure S11. Changes in expression levels of thyroid hormone-related genes in *Nkx2-1*$^{fl/fl}$ and *Nkx2-1*$^{\Delta T}$ mouse thyroids with combined exposure to iodine deficiency and radiation.** (A) Heatmap of the thyroid hormone-related genes generated with the RNA-seq data from *Nkx2-1*$^{fl/fl}$ and *Nkx2-1*$^{\Delta T}$ thyroids. A red arrow indicates the gene with significant up-regulation in *Nkx2-1*$^{\Delta T}$ thyroids. (B and C) Schematic representation of the time course (Left) and qRT-PCR for indicated thyroid hormone-related genes using the cDNAs from *Nkx2-1*$^{fl/fl}$ and *Nkx2-1*$^{\Delta T}$ thyroids used in Figure 3D for (B) and Figure 3H for (C) (Right). Data are presented as mean ± SD from individual mouse thyroid samples. For each gene, the mean value of *Nkx2-1*$^{fl/fl}$ thyroid samples was set as 1. *p<0.05, **p<0.01, ***p<0.005 by Student's *t*-tests.

**Table S1. Diffuse and focal hyperplasia incidence developed in *Nkx2-1*$^{fl/fl}$ and *Nkx2-1*$^{\Delta T}$ mouse thyroids under iodine deficiency and radiation exposure examined over 1 year after diet replacement.**

| Genotype | Diet/ Radiation | Diffuse hyperplasia (%) (mouse number) | Focal hyperplasia (%) (mouse number) | Total mouse number (N) |
|---|---|---|---|---|
| *Nkx2-1*$^{fl/fl}$ | CD | 0 (0) | 0 (0) | 21 |
| *Nkx2-1*$^{\Delta T}$ | | 0 (0) | 22* (5) | 23 |
| *Nkx2-1*$^{fl/fl}$ | CD+RAD | 0 (0) | 5 (1) | 22 |
| *Nkx2-1*$^{\Delta T}$ | | 0 (0) | 21 (6) | 28 |
| *Nkx2-1*$^{fl/fl}$ | IDD | 83 (15) | 50 (9) | 18 |
| *Nkx2-1*$^{\Delta T}$ | | 87 (26) | 73 (22) | 30 |
| *Nkx2-1*$^{fl/fl}$ | IDD+RAD | 74 (20) | 56 (15) | 27 |
| *Nkx2-1*$^{\Delta T}$ | | 74 (25) | 68 (23) | 34 |

Frequency of diffuse and focal hyperplasia developed in *Nkx2-1*$^{fl/fl}$ and *Nkx2-1*$^{\Delta T}$ mice used in Table 1 is summarized. Note that one *Nkx2-1*$^{\Delta T}$ mouse sample in IDD+RAD group was occupied with cancer tissue and this sample was removed from evaluation and is not included in the total mouse number.

* $p<0.05$ vs *Nkx2-1*$^{fl/fl}$ in CD group by Fisher's exact test calculated as a two-group comparison.

**Table S2. Effects of heterozygous and homozygous deletion of *Nkx2-1* on thyroid adenoma and carcinoma incidence in mice exposed to a combination of iodine deficiency and radiation.**

| Genotype | Diet/ Radiation | Adenoma(%)* (mouse number) | Carcinoma (%) (mouse number) | Total mouse number (N) |
|---|---|---|---|---|
| *Nkx2-1*$^{fl/wt}$ or *Nkx2-1*$^{fl/fl}$ | IDD+RAD | 25 (5) | 5 (1) | 20 |
| *Nkx2-1*$^{H\Delta T}$ | | 18 (2) | 0 (0) | 11 |
| *Nkx2-1*$^{\Delta T}$ | | 63 (10) | 13 (2) | 16 |

Frequency of adenoma and carcinoma developed in *Nkx2-1*$^{fl/wt}$ or *Nkx2-1*$^{fl/fl}$, *Nkx2-1*$^{H\Delta T}$ and *Nkx2-1*$^{\Delta T}$ mice exposed to a combination of iodine deficiency and radiation. See details in Materials & Methods. *$p<0.05$ by the Chi-square test for multiple group comparison.

**Table S3. Significantly enriched pathways in *Nkx2-1*$^{\Delta T}$ mouse thyroids using WikiPathways subset.**

| NAME | SIZE | ES | NES | NOM p-val | FDR q-val |
|---|---|---|---|---|---|
| WP_METAPATHWAY_BIOTRANSFORMATION | 91 | -0.662 | -1.860 | 0.000 | 0.036 |
| WP_SPLICING_FACTOR_NOVA_REGULATED_SYNAPTIC_PROTEINS | 38 | -0.657 | -1.770 | 0.000 | 0.075 |
| WP_TCA_CYCLE | 30 | -0.664 | -1.661 | 0.000 | 0.159 |
| WP_OXIDATIVE_STRESS_RESPONSE | 26 | -0.580 | -1.643 | 0.014 | 0.147 |
| WP_AMINO_ACID_METABOLISM | 79 | -0.638 | -1.637 | 0.000 | 0.132 |
| WP_P38_MAPK_SIGNALING_PATHWAY | 32 | -0.455 | -1.618 | 0.000 | 0.141 |
| WP_CHOLESTEROL_METABOLISM (Bloch and Kandutsch-Russel pathways) | 47 | -0.720 | -1.590 | 0.022 | 0.162 |
| WP_MITOCHONDRIAL_LONG_CHAIN_FATTY_ACID_BETAOXIDATION | 16 | -0.710 | -1.577 | 0.000 | 0.157 |
| WP_GLUTATHIONE_METABOLISM | 18 | -0.646 | -1.577 | 0.012 | 0.141 |
| WP_MAPK_CASCADE | 28 | -0.410 | -1.551 | 0.000 | 0.175 |
| WP_ONECARBON_METABOLISM_AND_RELATED_PATHWAYS | 43 | -0.596 | -1.540 | 0.000 | 0.178 |
| WP_OXIDATIVE_STRESS_AND_REDOX_PATHWAY | 78 | -0.524 | -1.502 | 0.029 | 0.247 |

Significantly enriched pathways from the WikiPathways Canonical subset (FDR q-value<0.25) in *Nkx2-1*$^{\Delta T}$ thyroids, ranked by normalized enrichment score (NES). Oxidative stress-related pathways are highlighted in yellow.

**Table S4. Significantly enriched pathways in *Nkx2-1*$^{\Delta T}$ mouse thyroids using MH mouse hallmark gene sets.**

| NAME | SIZE | ES | NES | NOM p-val | FDR q-val |
|---|---|---|---|---|---|
| HALLMARK_PROTEIN_SECRETION | 94 | -0.565 | -1.799 | 0.000 | 0.097 |
| HALLMARK_PEROXISOME | 92 | -0.510 | -1.736 | 0.000 | 0.110 |
| HALLMARK_FATTY_ACID_METABOLISM | 139 | -0.558 | -1.695 | 0.000 | 0.098 |
| HALLMARK_MTORC1_SIGNALING | 194 | -0.513 | -1.677 | 0.000 | 0.092 |
| HALLMARK_MYC_TARGETS_V2 | 58 | -0.541 | -1.631 | 0.035 | 0.107 |
| HALLMARK_GLYCOLYSIS | 181 | -0.433 | -1.617 | 0.009 | 0.099 |
| HALLMARK_ADIPOGENESIS | 199 | -0.538 | -1.613 | 0.078 | 0.089 |
| HALLMARK_OXIDATIVE_PHOSPHORYLATION | 193 | -0.462 | -1.597 | 0.095 | 0.092 |
| HALLMARK_CHOLESTEROL_HOMEOSTASIS | 68 | -0.701 | -1.585 | 0.000 | 0.093 |
| HALLMARK_MYC_TARGETS_V1 | 193 | -0.352 | -1.568 | 0.189 | 0.100 |
| HALLMARK_BILE_ACID_METABOLISM | 86 | -0.465 | -1.557 | 0.018 | 0.096 |
| HALLMARK_UNFOLDED_PROTEIN_RESPONSE | 108 | -0.381 | -1.526 | 0.079 | 0.115 |
| HALLMARK_ESTROGEN_RESPONSE_LATE | 172 | -0.515 | -1.474 | 0.006 | 0.162 |
| HALLMARK_XENOBIOTIC_METABOLISM | 157 | -0.481 | -1.466 | 0.011 | 0.156 |
| HALLMARK_ESTROGEN_RESPONSE_EARLY | 184 | -0.500 | -1.464 | 0.061 | 0.146 |
| HALLMARK_NOTCH_SIGNALING | 30 | -0.458 | -1.405 | 0.090 | 0.204 |
| HALLMARK_SPERMATOGENESIS | 72 | -0.530 | -1.395 | 0.109 | 0.207 |
| HALLMARK_P53_PATHWAY | 186 | -0.336 | -1.374 | 0.040 | 0.218 |
| HALLMARK_KRAS_SIGNALING_DN | 109 | -0.608 | -1.364 | 0.035 | 0.218 |
| HALLMARK_REACTIVE_OXYGEN_SPECIES_PATHWAY | 47 | -0.299 | -1.336 | 0.141 | 0.242 |
| HALLMARK_HEDGEHOG_SIGNALING | 33 | -0.523 | -1.328 | 0.112 | 0.239 |

Significantly enriched MH pathways (FDR q-value<0.25) in *Nkx2-1*$^{\Delta T}$ thyroids, ranked by NES. Oxidative stress-related pathways are highlighted in yellow.

**Table S5. qRT-PCR primers used in this study.**

Mouse qRT-PCR primers

| Gene | Forward (5'-3') | Reverse (3'-5') | References |
|---|---|---|---|
| *Hprt1* | CTGGTTAAGCAGTACAGCCCCA | GGTCCTTTTCACCAGCAAGCT | Shirakawa et al. [1]; Shirai et al. [2] |
| *Actb* | ACACCCGCCACCAGTTC | TACAGCCCGGGGAGCAT | Mehta et al. [3] |
| *Rps18* | CAGCCAGGTTCTGGCCAACGG | ATACACCCACAGTTCGGCCCCTG | Mu et al. [4] |
| *Nkx2-1* (#1) | ACAGCCAAGCAAATTCAACC | GGGTGCATCCACAGAAAAGT | Shirai et al. [2] |
| *Nkx2-1* (#2) | AGCACACGACTCCGTTCTC | GCCCACTTTCTTGTAGCTTTCC | Gotoh et al. [5]; Shirai et al. [2] |
| *Sod1* | CAGAAGGCAAGCGGTGAAC | CAGCCTTGTGTATTGTCCCCATA | Kim et al. [6] |
| *Sod2* | TACAACTCAGGTCGCTCTTCAGC | AGCCTCCAGCAACTCTCCTTT | Sidarala et al. [7] |
| *Sod3* | CCCACCCCCAAGTTCCAT | AAAGGTTCCCAAATACTCTCTCTAAGG | Kim et al. [6] |
| *Cat* | GGACGCTCAGCTTTTCATTC | TTGTCCAGAAGAGCCTGGAT | Kim et al. [6] |
| *Gpx1* | CGCTCTTTACCTTCCTGCGGAA | AGTTCCAGGCAATGTCGTTGCG | Kang et al. [8] |
| *Gpx2* | ATCAAACGGCTCCTCAAAGT | GGGACGATATTCAGGGAATG | Carlson et al. [9] |
| *Gpx3* | GATGTGAACGGGGAGAAAGA | TTCATGGGTTCCCAAAAGAG | Kim et al. [6] |
| *Gpx4* | TAAGAACGGCTGCGTGGT | GTAGGGGCACACACTTGTAGG | Kim et al. [6] |
| *Gpx7* | CGACTTCAAGGCGGTCAACATC | AAGGCTCGGTAGTTCTGGTCTG | Kang et al. [8] |
| *Gpx8* | ACATTCCCCATCTTCCACAA | ATTCCACCTTGGCTCCTTCT | Pineiro-Hermida et al. [10] |
| *Prdx1* | ACACCCAAGAAACAAGGAGGATT | CAACGGGAAGATCGTTTATTGTTA | Fatma et al. [11] |
| *Prdx2* | AACGCGCAAATCGGAAAGT | AGTCCTCAGCATGGTCGCTAA | Fatma et al. [11] |
| *Prdx3* | GGCCACATGAACATCACACTGT | CAAACTGGAACGCCTTTACCA | Fatma et al. [11] |
| *Prdx4* | TCCTGTTGCGGACCGAAT | GATCTTGGCTTTGCTTAGATGCA | Wang et al. [12] |
| *Prdx5* | GAAAGAAGCAGGTTGGGAGTGT | CCCAGGGACTCCAAACAAAA | Fatma et al. [11] |
| *Prdx6* | TTCAATAGACAGTGTTGAGGATCA | CGTGGGTGTTTCACCATTG | Chhunchha et al. [13] |
| *Gstm2* | TACCTTGCCCGAAAGCACAA | ATCTTCTCAGGGAGACCCTCT | Li et al. [14] |
| *Gstm5* | TCATCCAAGTCTATGGTTCTGGG | CCACAGATGTACCGTTTCTCCT | Hou et al. [15] |
| *Gstm7* | CTTATGGACAACCGCATGGTG | TACCCTGGCTTCAGCTTCTCA | Yip et al. [16] |
| *Duox1* | ACCAGAACATTGCGATGTATGAG | AGAAATGGACGGTATCCTGGA | Schiffers et al. [17] |
| *Duox2* | GGACAGCATGCTTCCAACAAGT | GCCTGATAAACACCGTCAGCA | Grasberger et al. [18]; Shirai et al. [2] |
| *Duoxa1* | CCCACAGGATGCAGCCTCAC | ACCGGTAGTGGGGGCTCAAG | Cheon et al. [19] |
| *Duoxa2* | CGTTAACATTACACTCCGAGGAAC | CAGAATGCCACCCACAGTGT | Grasberger et al. [18] |
| *Slc5a5* | GGTGCTCTCATCAGCTACCT | GTCGCAGCAGGGATGTCT | Grasberger et al. [18], Shirai et al. [2] |
| *Tshr* | TCTCAAAAAGCTCCCGCTGT | AGACTCCAGGATTCCCCTGAT | Shirai et al. [2] |
| *Tpo* | GCAGGTGGACACATGCTGA | GTCTGGCTCCAAAGCAGTGA | Grasberger et al.[18]; Shirai et al.[2] |
| *Tg* | TCCGGAGGAAAGTTGTGCTG | GCCGCTCACACTCAAAGAAC | van der Vaart et al.[20]; Shirai et al. [2] |
| *Acox1* | GGATGGTAGTCCGGAGAACA | AGTCTGGATCGTTCAGAATCAAG | He et al. [21] |
| *Acox2* | CCTTCCTAGACCTGCTTCCC | TGTCCGTCATAACAGCCAAG | He et al. [21] |
| *Acox3* | CTTCTGAGAAACGGGGACAA | GCTCGGTAGGCACTAAGAGG | He et al. [21] |

qRT-PCR primers to detect overexpressed rat *Nkx2-1* in 458s- or 459-TNKX cells

| Gene | Forward (5'-3') | Reverse (3'-5') | References |
|---|---|---|---|
| *Nkx2-1* | AGCACACGACTCCGTTCTC | GCCCACTTTCTTGTAGCTTTCC | Gotoh et al. [5]; Shirai et al. [2] |

**Table S6. Antibodies used in this study.**

| Antibody | Catalogue number | Company |
|---|---|---|
| Rabbit monoclonal anti-TTF1/Nkx2-1 | ab76013 | abcam |
| Mouse monoclonal anti-SOD1 | sc-271014 | Santa Cruz Biotechnology |
| Mouse monoclonal anti-SOD2 | sc-137254 | Santa Cruz Biotechnology |
| Mouse monoclonal anti-SOD3 | sc-271170 | Santa Cruz Biotechnology |
| Rabbit polyclonal anti-GPX2 | ab137431 | abcam |
| Rabbit polyclonal anti-Catalase (CAT) | ab52477 | abcam |
| Rabbit monoclonal anti-Catalase (CAT) | #14097 | Cell Signaling Technology |
| Rabbit polyclonal anti-PRDX6 | ab133348 | abcam |
| Rabbit monoclonal anti-Phospho-Histone H2A.X (Ser139) | #9718 | Cell Signaling Technology |
| Mouse monoclonal anti-Phospho-Histone H2A.X (Ser139) | #80312 | Cell Signaling Technology |
| Mouse monoclonal anti-Histone H2A.X | sc-517336 | Santa Cruz Biotechnology |
| Rabbit polyclonal anti-Vinculin | #4650 | Cell Signaling Technology |
| Rabbit monoclonal anti-GAPDH | #2118 | Cell Signaling Technology |
| Mouse monoclonal anti-$\beta$-Actin | sc-47778 | Santa Cruz Biotechnology |
| Mouse monoclonal anti-$\alpha$-Tubulin | sc-32293 | Santa Cruz Biotechnology |
| Goat anti-rabbit IgG, HRP-linked antibody | #7074 | Cell Signaling Technology |
| Horse anti-mouse IgG, HRP-linked antibody | #7076 | Cell Signaling Technology |
